# A self-regenerating synthetic cell model

**DOI:** 10.1101/2020.07.03.185900

**Authors:** Barbora Lavickova, Nadanai Laohakunakorn, Sebastian J. Maerkl

**Affiliations:** Institute of Bioengineering, School of Engineering, École Polytechnique Fédérale de Lausanne, Lausanne, Switzerland; Institute of Quantitative Biology, Biochemistry, and Biotechnology, School of Biological Sciences, University of Edinburgh, Edinburgh, United Kingdom

## Abstract

Self-regeneration is a fundamental function of all living systems. Here we demonstrate molecular self-regeneration in a synthetic cell model. By implementing a minimal transcription-translation system within microfluidic reactors, the system was able to regenerate essential protein components from DNA templates and sustained synthesis activity for over a day. By mapping genotype-phenotype landscapes combined with computational modeling we found that minimizing resource competition and optimizing resource allocation are both critically important for achieving robust system function. With this understanding, we achieved simultaneous regeneration of multiple proteins by determining the required DNA ratios necessary for sustained self-regeneration. This work introduces a conceptual and experimental framework for the development of a self-replicating synthetic cell.

## Introduction

Bottom-up construction of a self-replicating synthetic cell that exhibits all the hallmarks of a natural living system is an outstanding challenge in synthetic biology [1, 2, 3]. While this goal is ambitious, progress is rapidly accelerating, and key structures and functions required for constructing a synthetic cell, including compartmentalization [4, 5, 6], mobility and shape [7, 8, 9], metabolism [10, 11], communication [12, 13], and DNA replication [14, 15], have recently been demonstrated, suggesting that integration of these subsystems into a functional synthetic cell may be an attainable goal.

A biochemical system able to fully self-regenerate or self-replicate, is a crucial requirement for construction of a synthetic cell. A self-replicating artificial system has been first proposed by von Neumann in the 1940s [16]. Von Neumann developed the concept of a universal constructor, which is an abstract machine capable of self-replication using a set of instructions, external building blocks, and energy. So far, universal constructors have only been implemented *in silico* in the form of cellular automata [17]. Similar concepts have been explored experimentally with auto-catalytic chemical systems [18] and self-replicating ribozymes [19]. A self-replicating biochemical system is strictly analogous to the universal constructor in that it would be capable of self-replication using instructions encoded in DNA while being supplied with building blocks and energy (Fig. 1A). A physical implementation of a universal constructor could therefore be theoretically achieved by a minimal recombinant transcription-translation system capable of regenerating all of its components including proteins, ribosomes, tRNAs, and DNA [20]. DNA replication has recently been demonstrated in *vitro* [14, 15] and progress is being made in reconstituting ribosomes [21, 22]. Here we demonstrate the principle steps towards constructing a universal biochemical constructor by creating a system capable of sustained self-regeneration of proteins essential for transcription and translation.

**Figure 1:**
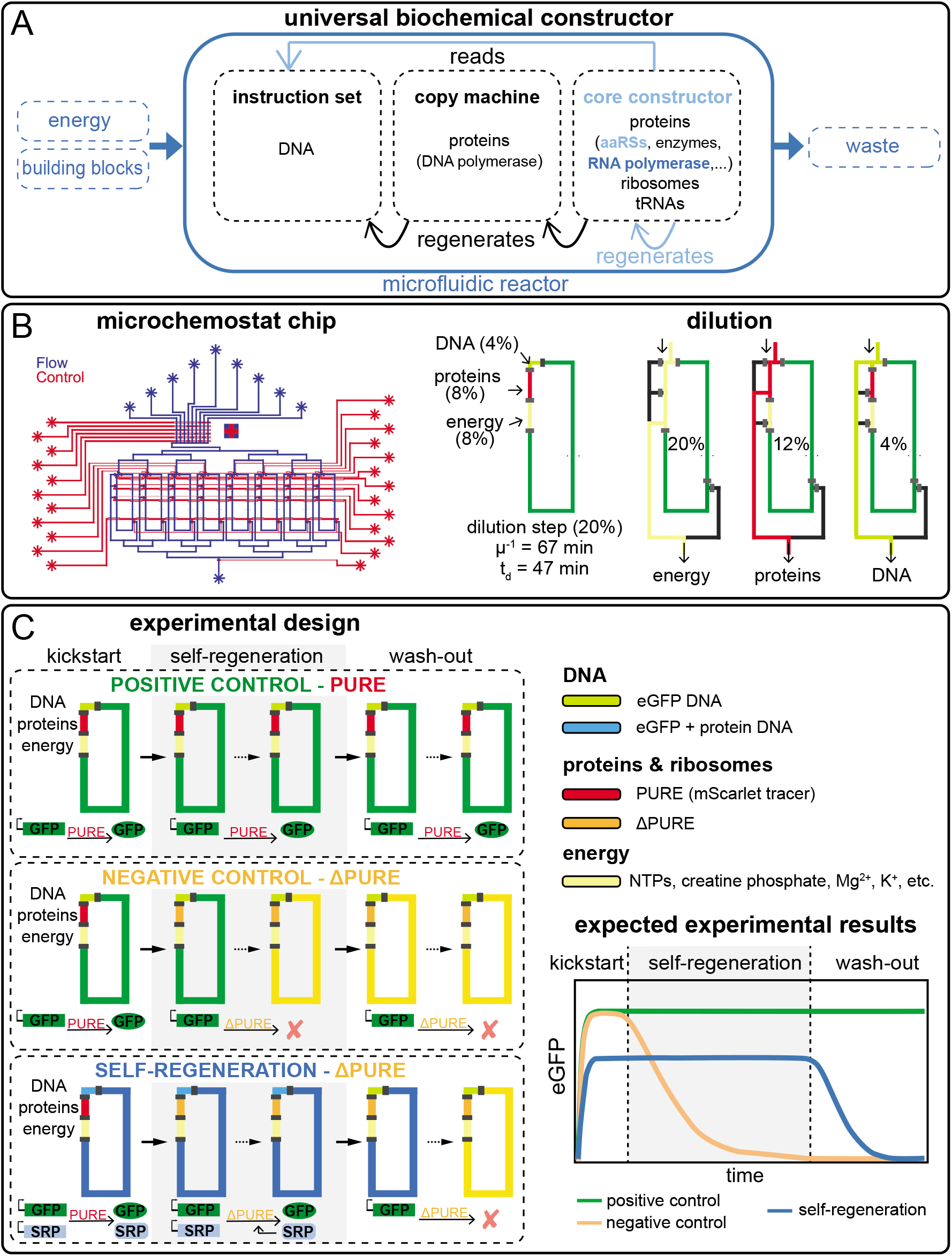
**(A)** Diagram of the universal biochemical constructor concept. Systems, components, and functions colored in blue and light-blue were fully or partially implemented in this work, respectively. **(B)** Design schematic of the microfluidic device with eight individual chemostat reactors. Design and functional details are provided in Supplementary Fig. S1. A schematic representation of one dilution cycle were 20% reaction volume is replaced every 15 min. Dilution rate *μ* = −ln(C_t_/C_0_)·t^−1^, residence time *μ*^−1^ and dilution time t_d_ = ln(2)·*μ*^−1^. One dilution cycle consists of three steps: energy solution is loaded via the 20% segment, protein and ribosome solution is flushed through the 12% segment, and DNA solution through the 4% segment, resulting in the desired composition of 8%, 8%, and 4 %, respectively. **(C)** Experimental design, including the three experiment phases: kickstart, self-regeneration, and wash-out. A schematic showing the expected results for the different experimental phases indicating early cessation of synthesis activity for the negative control, continuous synthesis activity in the positive control, and continuous synthesis during the self-regeneration phase followed by cessation of synthesis activity during the wash-out phase.

Development of a transcription-translation system capable of self-regeneration faces several challenges. First, synthesis capacity of the system in terms of its protein synthesis rate must be sufficient to regenerate the necessary components. This problem is exacerbated by the fact that protein synthesis capacity drastically decreases in a non-optimal system [23, 24, 25]. Second, the components being regenerated must be functionally synthesized which may require chaperones, and modifying enzymes. And third, the reaction must take place in an environment that allows continuous and sustained regeneration.

Here we employ continuous transcription-translation reactions operating inside microfluidic reactors [26] to demonstrate self-regeneration of essential protein components. We chose the PURE (protein synthesis using recombinant elements) system [27] as a viable starting point for achieving self-regeneration because of its minimal nature as well as its defined and adjustable composition [28]. Batch expression experiments combined with polyacrylamide gel electrophoresis (PAGE) and mass-spectrometric (MS) analysis indicated that the PURE system should be able to synthesize around 70% of all *E. coli* proteins [29]. Moreover, it was recently shown that co-expression of multiple PURE components in a single batch reaction yielded the required concentrations for self-replication [15]. However, these experiments didn’t determine whether proteins were functionally synthesized, which varies largely for proteins expressed in the PURE system [30, 31]. Two other studies showed that the 30S ribosomal subunit [22], and nineteen of twenty aaRSs, can be functionally synthesized in the PURE system [32]. All of those experiments were performed in batch or continuous-exchange formats and self-regeneration of any component has yet to be demonstrated.

Our approach using the PURE system, microfluidic chemostats, and monitoring fluorescent protein production, allows activity and performance of self-regeneration to be assessed in real-time. We implemented a ‘kick-start’ method to ‘boot-up’ regeneration of essential PURE proteins from DNA templates. We demonstrate the concept and feasibility of this approach by regenerating different aminoacyl-tRNA synthetases (aaRSs). We also regenerated T7 RNA polymerase (RNAP) and mapped system optimality by varying T7 RNAP DNA concentration and were able to explain the observed genotype-phenotype landscape with a biophysical resource limitation model. We go on to show that several proteins can be regenerated simultaneously by regenerating up to seven aaRSs. This proof-of-principle work demonstrates the first steps towards constructing a self-replicating transcription-translation system and provides a viable approach for developing and optimizing other critical sub-systems including DNA replication, ribosome synthesis, and tRNA synthesis, with the goal of achieving a self-replicating biochemical constructor in the near term and ultimately a viable synthetic cell.

## Results

### Experimental design

To maintain continuous cell-free reactions we improved a microfluidic chemostat previously used for implementing and forward engineering genetic networks *in vitro* [26, 33]. The device consists of 8 independent, 15 nL reactors, with fluidically hard-coded dilution fractions defined by reactor geometry, as opposed to the original device which used peristaltic pumps for metering (Fig. 1B, Supplementary Fig. S1) [34]. During experiments 20% of the reactor volume was replaced every 15 min with a ratio of 2:2:1 for energy, protein/ribosome, and DNA solution, respectively, resulting in an effective dilution time of ~ 47 min (Fig. 1A, Supplementary Table S1, S2). Another key improvement was the supply of multiple solutions without the need for cooling. This was achieved by storing the energy and protein components separately, which when stored pre-mixed and without cooling resulted in non-productive resource consumption [35]. Secondly, reaction temperature was set to 34°C, which decreased PURE degradation with only a minor decrease in protein synthesis rate (Supplementary Fig. S2). Lastly, as the redox reagent used in the PURE system is known to degrade rapidly, we eliminated 1,4-dithiothreitol (DTT) in the energy solution and instead added tris(2 carboxyethyl)phosphine (TCEP) to the energy and protein solutions. To allow PURE system modification and omission of protein components we produced our own PURE system based on the original formulation [36, 23]. For each protein regenerated, we produced a ΔPURE system lacking that particular protein or proteins. This allowed us to validate that the omitted protein is essential for system function. We furthermore adjusted PURE protein composition by reducing the concentration of several aaRSs (Supplementary Table S3, S4).

In all experiments we expressed a fluorescent protein (eGFP) as an indicator of functional self-regeneration and to provide a quantitative readout of protein synthesis capacity. We developed a ‘kick-start’ method to enable the system to self-regenerate proteins from DNA templates (Fig. 1C). The experimental design involves three distinct phases: kick-start, self-regeneration, and wash-out. The kick-start phase is required to allow a productive switch from a complete to a ΔPURE system to occur. The self-regeneration phase tests whether the system functionally regenerated the omitted protein component or components, and the washout phase serves as a control to prove that the omitted component or components were indeed essential for system function. In the kick-start phase, which lasts for the first 4h, linear DNA templates coding for eGFP and the protein to be regenerated are added to a complete PURE system. This leads to the expression of eGFP and the protein to be regenerated. In the self-regeneration phase, the full PURE is gradually replaced with a ΔPURE solution lacking the particular protein that is to be regenerated. Thus at steady state, the system will remain functional only through self-regeneration of the omitted protein. Finally, in the wash-out phase, DNA encoding the protein being regenerated is no longer added to the system leading to dilution of the protein being regenerated. Once a critical concentration for the regenerated protein is reached overall protein synthesis falls and ultimately ceases.

We implemented two additional control reactions in most experiments. Positive controls use full PURE and express only eGFP during all three phases and serve as a validation of steady-state chemostat function and a reference point for maximal protein synthesis capacity of an unloaded and optimal PURE reaction. Negative controls switch between complete and ΔPURE, but don’t contain DNA template for the omitted protein component. This confirms that without self-regeneration, protein synthesis activity is indeed rapidly lost. We spiked the full PURE protein fraction with an mScarlet tracer to confirm that all fluid exchanges take place and the device functioned correctly.

### Aminoacyl-tRNA synthetase regeneration

As a proof-of-concept and validation of the experimental design, we tested regeneration of two aaRSs: Asparaginyl-tRNA synthetase (AsnRS) and Leucyl-tRNA synthetase (LeuRS) (Fig. 2A). We first carried out batch experiments to ascertain synthesis of the synthetases in our PURE system (Supplementary Fig. S3). We also validated that both synthetases are essential by omitting them individually from a PURE reaction (Fig. 2B). When we used the original PURE system’s aaRS concentrations, decreases in protein synthesis activity were observed only after extended washout periods because the critical aaRS concentrations were reached only after numerous dilution cycles (data not shown). We therefore reduced the concentrations of the aaRSs being regenerated so that fast activity declines during wash-out occurred, while preserving high protein synthesis rates (Supplementary Fig. S4, Table S3).

**Figure 2:**
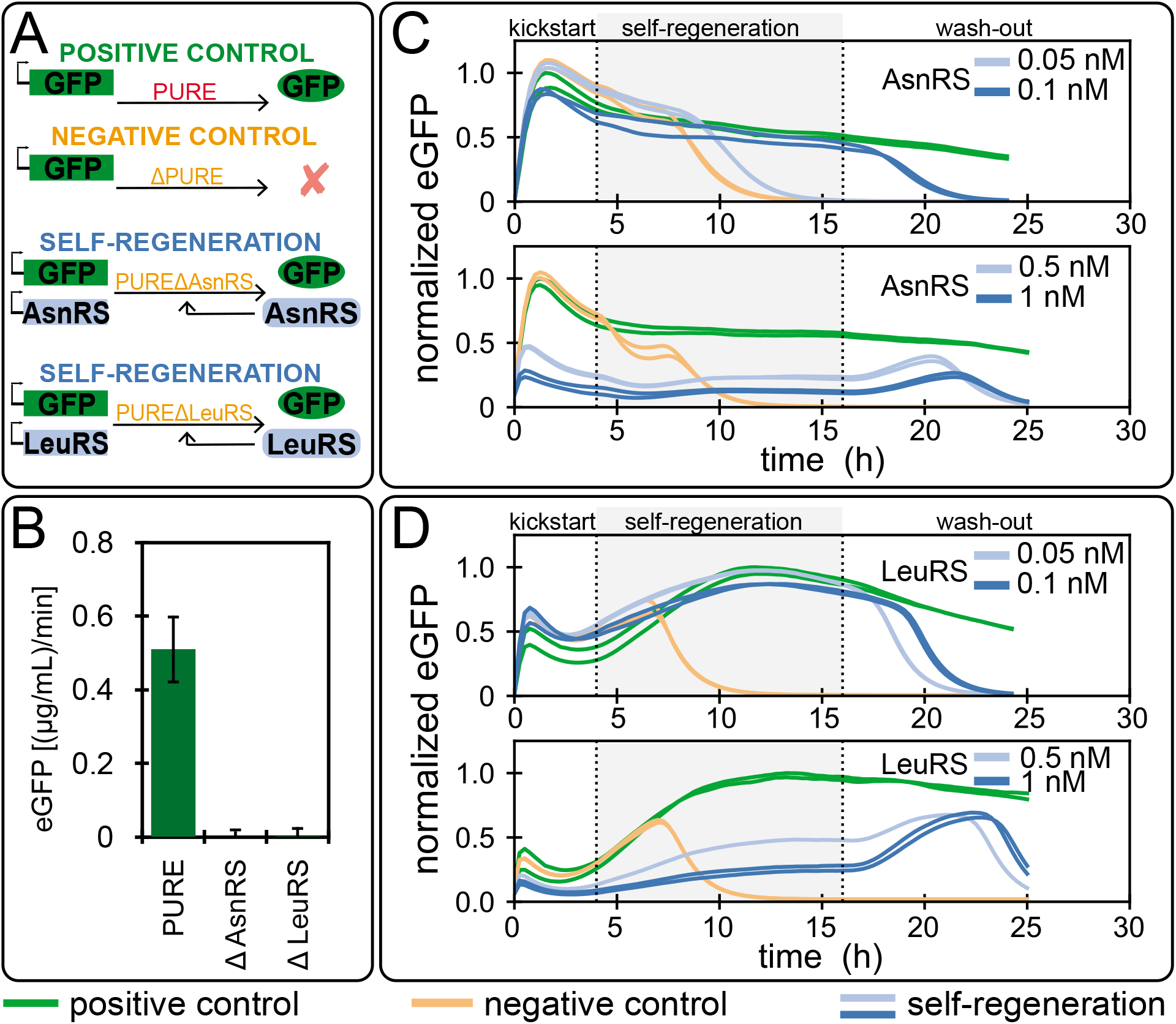
Aminoacyl-tRNA synthetase regeneration: **(A)** overview of the different aaRSs regeneration experiments. **(B)** eGFP batch synthesis rates for the full PURE system and AsnRS or LeuRS ΔPURE systems. Values are mean ± s.d. for PURE system (n=16), and mean ± 10x s.d. for ΔPURE systems (n=2). Self-regeneration experiments for **(C)** AsnRS **(D)** LeuRS at different DNA concentrations. Results for all DNA concentrations tested and corresponding mScarlet traces can be found in Supplementary Fig. S6. The level of eGFP is normalised to the maximum level attained in the positive control. The composition of PURE systems used are given in Supplementary Table S3, 2 nM of eGFP template was used for all experiments, aaRS DNA template concentrations are indicated.

We achieved successful self-regeneration for both AsnRS and LeuRS and complete loss of protein synthesis activity during wash-out (Fig. 2C, D). We tested four DNA concentrations for each aaRSs. AsnRS and LeuRS regeneration at DNA concentrations of 0.1 nM and 0.05 nM, respectively, resulted in high system activity comparable to the positive control throughout the self-regeneration phase. If an insufficient DNA template concentration of 0.05 nM was provided for AsnRS, a decrease in eGFP fluorescence was observed identical to the negative control but with a slight delay. A twofold difference in DNA template concentration thus resulted in either optimal self-regeneration or complete system failure. For LeuRS a similar two-fold change was less consequential with either concentration resulting in self-regeneration, but with slightly lower expression obtained for the higher concentration of 0.1 nM. Higher DNA concentrations resulted in robust but markedly lower system activity for both aaRSs. These studies showed that our experimental design enables self-regeneration and that self-regeneration can be achieved with two different aaRSs.

DNA input concentration is critically important for system function. When higher than optimal DNA concentrations were used, we observed successful and robust self-regeneration, as indicated by the maintenance of synthesis activity above negative control levels, but considerably lower eGFP expression levels as compared to the positive control. Because no negative effects were observed in batch reactions for high aaRS protein concentrations in the PURE system (Supplementary Fig.S4) [37], we attribute this effect to a resource competition or loading effect between the protein being regenerated and eGFP [38]. The onset of this loading effect can be estimated by measuring the DNA concentration for which system output saturates, which is ~ 1 nM for the PURE system (Supplementary Fig. S5). eGFP DNA template is present at a concentration of 2 nM in all experiments and is thus fully loading the system. Any additional DNA added to the system will thus give rise to resource competition effects.

A simple resource competition model gives rise to a couple of specific predictions. First, the level of eGFP synthesized during self-regeneration should never rise above the positive control, assuming that the concentration of the self-regenerated protein is at an optimal level in the positive control. This is because synthesis of an additional protein leads to resource competition and lower eGFP levels. Low concentrations of aaRS DNA has a minimal loading effect since the ratio of aaRS to eGFP DNA is small. As the concentration of aaRS DNA is increased the loading effect becomes stronger, leading to a noticeable decrease in eGFP levels. The second prediction is that eGFP levels can exhibit a transient peak during washout phase. This occurs because loading decreases before the regenerated protein is diluted below critical levels. This is evident in our experiments with high load levels (high aaRS input DNA concentrations), where a transient spike in eGFP expression occurred during wash-out before a decrease was observed (Fig. 2C-D, Supplementary Fig. S6).

To approximate the optimal DNA input concentrations for self-regeneration, we estimated aaRS protein synthesis rates for different DNA concentrations by using the ratio of aaRS to eGFP DNA, while assuming the same synthesis rate for all proteins, and comparing them to the estimated synthesis rate required to reach the minimum concentration needed for each aaRS (Supplementary Fig. S7A, Supplementary Table S4). In agreement with the observed data, we estimated 0.1 and 0.05 nM of DNA for AsnRS and LeuRS, respectively. Moreover, we confirmed these estimates based on the drop in eGFP synthesis rate for different DNA input concentrations (Supplementary Fig. S7B).

### T7 RNAP regeneration

After testing two proteins essential for translation, we tested self-regeneration of an essential protein for transcription (Fig. 3A). For transcription the PURE system utilises T7 RNA polymerase (RNAP), a single 99 kDa protein. As before, we carried out batch experiments to validate T7 RNAP synthesis in the PURE system (Supplementary Fig. S3), and essentiality of T7 RNAP (Fig. 3B). T7 RNAP could be successfully regenerated in the system and we carried out extensive DNA template titrations with concentrations varying over three orders of magnitude (Fig. 3C, Supplementary Fig. S8). By omitting the wash-out phase and extending the self-regeneration phase to 26 hours, we showed that T7 RNAP can be regenerated at steady-state for over 25 hours with a DNA input concentration of 0.5 nM (Fig. 3D).

**Figure 3:**
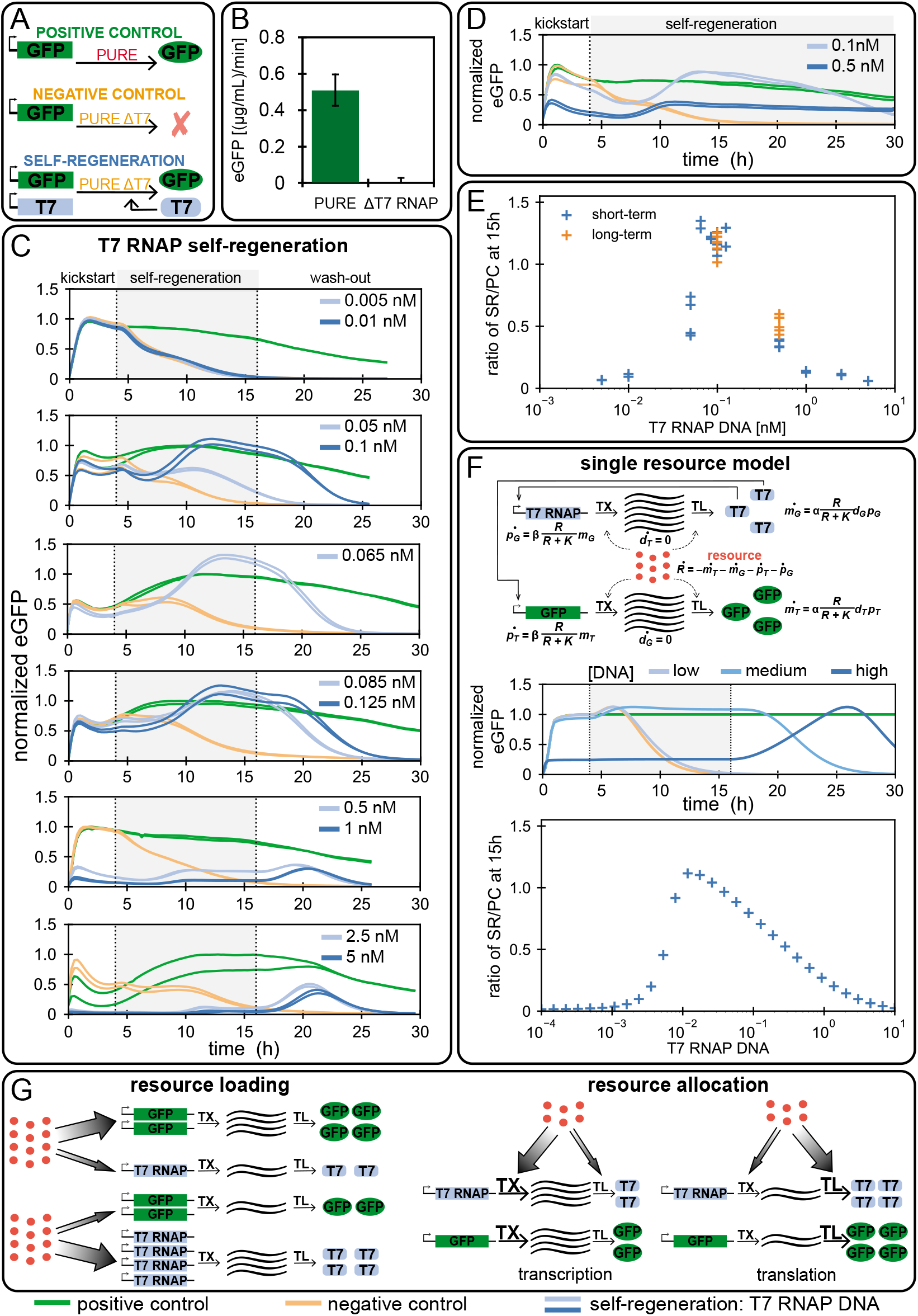
T7 RNAP self-regeneration: **(A)** Overview of the T7 RNAP regeneration experiment. **(B)** eGFP batch synthesis rates for the full PURE and T7 RNAP ΔPURE systems. Values are mean ± s.d. for the PURE system (n=10), and mean ± 10×s.d. for ΔPURE systems (n=2). **(C)** T7 RNAP regeneration at different DNA template concentrations. **(D)** Long-term regeneration experiment: the self-regeneration phase was extended by omitting the wash-out phase. The results for all DNA concentrations tested and the appropriate mScarlet traces can be found in Supplementary Fig. S8. The level of eGFP is normalised to the maximum level attained in the positive control experiments. The composition of the PURE system used for the self-regeneration experiments is given in Supplementary Table S3, 2 nM of eGFP template was used for all experiments, T7 RNAP DNA template concentrations are indicated. **(E)** Ratio of eGFP levels of the self-regeneration experiments and the positive control at 15 hours as a function of T7 RNAP DNA template concentration. Each data point represents a single measurement. **(F)** Our single resource model consists of seven ODEs and three parameters. DNA, mRNA, and protein concentrations are denoted by *d, m*, and *p*, and the subscripts *T* and *G* refer to T7 RNAP and eGFP, respectively. Simulation of a self-regeneration experiment: the switch between stages occurs at 4 and 16 hours. DNA for T7 RNAP was present at three qualitatively different concentrations, indicated as ‘low’, ‘medium’, and ‘high’. All concentrations are non-dimensional. The level of GFP is normalised by the maximum level attained in the positive control experiment. The negative control corresponds to *d_T_* = 0. **(G)** Schematic description of the concepts of resource loading and resource allocation. Resource loading is the distribution of a limited resource between two genes. Resource allocation is the distribution of a limited resource between transcription and translation.

To summarize the DNA titration results, we plotted eGFP expression levels as a function of T7 RNAP DNA template concentration at 11 hours of regeneration (corresponding to 15 hours after the start of the experiment), normalised to the positive control expression levels (Fig. 3E). For lower DNA concentrations (<0.05 nM) we observe little or no eGFP expression, which we attribute to insufficient synthesis of T7 RNAP, similar to the results obtained for aaRSs. For high T7 RNAP DNA template concentrations (≥ 1 nM) resource competition similar to what was observed for AsnRS and LeuRS was taking place. This is also supported by the observed peak during the wash-out phase for high input DNA template concentrations. Near optimal system performance within 80% of the control reaction occurred in a narrow DNA template concentration range of 0.65 nM to 0.125 nM. The curve is asymmetric, with higher sensitivity to low concentrations than to higher concentrations, providing insights into how system robustness can be engineered. Surprisingly, and unlike the aaRS experiments, we observed an expression maximum that rises to a level of 1.3 above the positive control reactions, indicating that a simple resource competition model cannot account for the observed behaviour.

To investigate whether our hypothesis of resource competition could be extended to explain the T7 RNAP observations, we created a minimal model of the transcription-translation system. While transcription-translation systems can be described at varying levels of granularity e.g. [39, 40, 41, 42], we chose to model the processes at the most coarse-grained level using coupled ordinary differential equations (ODEs) (Fig. 3F, Supplementary Fig. S9–S18, Supplementary Table S5).

The model consists of transcription and translation of eGFP and T7 RNAP, which consumes a single resource species *R*. This species is a lumped representation of CTP, UTP, ATP, GTP, and aminoacyl-tRNAs, which are consumed during transcription and/or translation. We model the transcription rate by a parameter *α*, linearly dependent on DNA and T7 RNAP concentration, and modulated by the availability of resources using a Hill function *R*/(*R* + *K*). Likewise, translation proceeds at a rate *β*, is linearly dependent on mRNA concentration, and is modulated by the same Hill function. The rate of consumption of *R* is equal to the summed transcription and translation rates. The complete model consists of seven ODEs and three parameters, and is solved between discrete dilution steps to simulate chemostat operation.

This minimal model successfully captures the observed qualitative behaviour including: 1) eGFP washout at low T7 RNAP DNA concentrations (*d_T_*) in the self-regeneration phase, 2) low eGFP production followed by a peak in the washout stage at high *d_T_*, and importantly 3) eGFP production in excess of the positive control at medium *d_T_* in the self-regeneration phase (Fig. 3F). At low *d_T_*, mRNA concentration is low, while resources are abundant; translation rate is thus limited by mRNA concentration. High *d_T_* leads to increased resource consumption, so despite the presence of large amounts of mRNA, translation is limited by resources. Further analysis reveals that at intermediate *d_T_* concentrations, eGFP production can increase above the positive control during the self-regeneration phase. The model predicts that this is due to a reallocation of resources from transcription to translation, once self-regeneration of T7 RNAP begins. This effect requires a resource-limited condition (Fig. 3G).

We developed an alternative resource-independent model which only takes into account translational loading through a shared translational enzyme, which can also capture the observed optimum in the SR/PC ratio (Supplementary Fig. S16). In this case, the optimum is due to a trade-off between mRNA concentration and enzyme availability. However it fails to predict the increase in eGFP production above the positive control during self-regeneration.

The modeling studies indicate that the requirement for an optimum in the SR/PC ratio is a coupling between eGFP and T7 RNAP expression, whether through a shared resource or a shared enzyme. However, the increase of eGFP above the positive control during the self-regeneration phase requires a resource-limited condition, and resource reallocation from transcription to translation (Fig. 3G). While both models can be combined, or extended to incorporate more realistic effects, such as saturation of transcription rates with substrate concentration, time delays in the various processes, and more intricate mechanisms of resource usage, none of these are required to explain our observations, apart from the essential feature of gene expression coupled through a shared resource.

### Regeneration of multiple components

Having demonstrated that proteins essential for translation or transcription could be regenerated individually, we explored whether multiple proteins could be regenerated simultaneously. We first tested if T7 RNAP, AsnRS, and LeuRS could be regenerated together. Initial DNA concentrations tested were 1× and 2× the minimal DNA concentrations which led to successful self-regeneration of individual proteins, but these concentrations were not sufficient for sustained self-regeneration of multiple proteins (Supplementary Fig. S19). Increasing DNA concentrations and maintaining 1:1 DNA template concentration ratios ultimately led to successful regeneration lasting 20-25 hours (Fig. 4A, Supplementary Fig. S19). Despite successfully regenerating for many hours, protein synthesis ultimately ceased under these conditions. Based on the T7 RNAP results and our computational modeling we hypothesized that a more optimal DNA ratio between T7 RNAP and the aaRSs needed to be established, as we previously observed strong resource loading effects by T7 RNAP and an apparent insensitivity of optimal T7 RNAP DNA concentration in respect to overall loading. Consequently, we decided to retain a relatively high DNA concentration of 0.5 nM for both aaRSs, and titrated T7 RNAP DNA template (Fig. 4A). This had the desired effect and resulted in sustained regeneration at a T7 RNAP DNA concentration of 0.2 nM.

**Figure 4:**
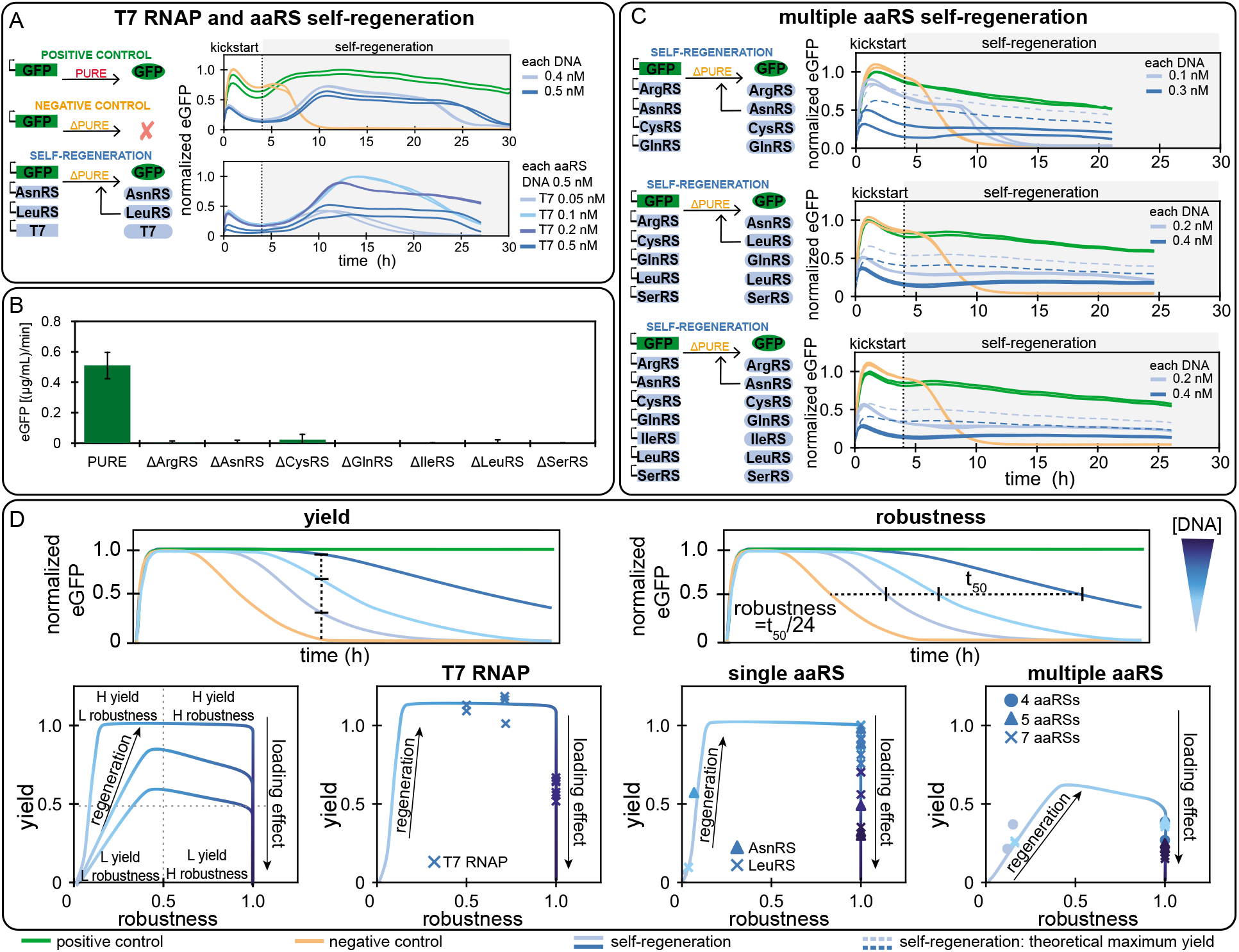
**(A)** Combined T7 RNAP, AsnRS, and LeuRS regeneration. Overview of the experiments on the left, with experimental results shown on the right. The top graph shows results for all DNA templates at concentrations of 0.4 or 0.5 nM. The bottom graph shows results for a titration of T7 RNAP DNA template with both aaRS DNA templates held constant at 0.5 nM. **(B)** eGFP batch synthesis rates for the PURE system and ΔPURE systems lacking additional aaRSs. Values are mean ± s.d. for PURE system (n=10), and mean ± 10×s.d. for ΔPURE systems (n=2). **(C)** Simultaneously regeneration of 4, 5, and 7 aaRSs. Overview of self-regeneration experiments on the left, with experimental results shown on the right. Results for all DNA concentrations tested and the corresponding mScarlet traces can be found in Supplementary Fig. S19. All eGFP traces were normalised to the maximum eGFP fluorescence output in the positive control, with exception of the T7 RNAP titration in panel **(A)** for which eGFP traces were normalised to the maximum eGFP fluorescence. PURE system compositions are given in Supplementary Table S3. 2 nM of eGFP DNA template was used in all experiments, other DNA template concentrations are indicated. **(D)** Schematic description of the definition of yield and robustness. Using these two terms we plotted theoretical curves for the relationship between DNA input concentration, yield, and robustness and superposed experimental values obtained from T7 RNAP, single aaRS, and multiple aaRS self-regeneration experiments. All three systems follow a similar trajectory and can be described in terms of Pareto optimality.

To explore the limits of the PURE transcription-translation system for self-regeneration, we tested whether several aaRSs could be regenerated simultaneously. We first carried out batch experiments to ensure efficient expression of the chosen aaRSs (Supplementary Fig. S3), as well as lack of expression if a given aaRS was omitted from the PURE system (Fig. 4B). As for the single aaRS experiments, we adjusted the concentrations of the various aaRS proteins (Supplementary Table S3) in the PURE system to ensure efficient wash-out. We gradually increased the number of aaRSs being regenerated from four to seven (Fig. 4C, Supplementary Fig. S20).

Based on eGFP synthesis rate and DNA ratios for the different conditions, we estimated that DNA inputs above 0.2 nM would be required for successful regeneration (Supplementary Fig. S21A). This was in agreement with the observed data and decreases in eGFP synthesis rate due to loading (Supplementary Fig. S21B). We observed successful self-regeneration of up to 22h for experiments with input DNA concentrations of 0.2 nM or above. DNA concentrations of 0.1 nM on the other hand led to rapid cessation of protein synthesis activity 10 hours into the experiment. Furthermore, when DNA input concentrations of 0.2 nM were used we saw variations in the length of self-regeneration amongst experiments (Supplementary Fig. S20). The estimated synthesis levels are much higher than the concentrations of most aaRS diluted out of the reactor each cycle, with exception of ArgRS, where 0.027 (*μ*g/mL)/min is diluted out, suggesting that optimization of DNA input for individual aaRSs could allow for better resource allocation and higher robustness.

eGFP levels are low when compared to the positive control. The positive control represents the maximum achievable eGFP steady-state levels in an otherwise unloaded system. Expression of 4-7 additional aaRS presents a considerable load on the system. When taking this load into account, self-regeneration of 4-7 aaRSs in addition to expressing eGFP reaches roughly 50% of the theoretically achievable yield (Fig. 4C), indicating that the total synthesis capacity of the system remained quite high.

These experimental results suggest that achieving successful self-regeneration depends on an interplay of several factors. To more quantitatively describe the system we defined the terms *yield* and *robustness* (Fig. 4D). We define *yield* as the level of non-essential protein such as eGFP that the system can synthesize during self-regeneration. In the case where an essential protein, for instance an aaRS, is missing, yield is zero. Expressing the aaRS will increase the yield, up to a point where the system’s resources are preferentially directed towards aaRS production. At that point system yield begins to decrease again due to loading. A second important parameter is system *robustness*. We consider a robust system to be able to sustain self-regeneration for at least 24 hours. A non-robust system may temporarily reach steady-state self-regeneration, but changes in synthesis rates, DNA concentrations, or environmental conditions, can cause it to cease functioning. We therefore define robustness as the time the system self-regenerates beyond the negative control, normalized by 24 hours. A system that self-regenerates for 24 hours or longer receives a robustness score of 1 while systems that cease regeneration before 24 hours receive a score between between 0 and 1.

Given these two parameters: yield and robustness, one can now describe the system in terms of Pareto optimality and determine whether there exists a trade-off between yield and robustness. In Figure 4D we show the calculated values of yield against robustness for our experimental observations, as well as the theoretically expected relationship of yield, robustness, and DNA concentration. For an essential protein, increasing its expression constrains the yield of the system onto a Pareto front [43]. This is because low expression of that protein leads to large increases in yield as the protein begins to confer its advantage on the system. Above a critical concentration, the system is able to continuously regenerate, corresponding to a robustness of 1. Expressing the protein at higher levels than the critical value incurs a cost on the system, which is exhibited by decreasing yield due to loading.

For single protein self-regeneration it is indeed possible to reach maximal yield and robustness. However, whether that situation can be attained or not depends on the activity of the essential protein. Proteins with low activity require higher concentrations, and hence more resources, to produce. This thus limits the attainable yield of the system, and shifts the yield-robustness curve downwards. In severe cases (which we do not observe), yield may never reach 1. Regeneration of multiple proteins falls into this category: when several essential proteins are being regenerated, the available capacity to express other proteins becomes less. Nonetheless, it is possible to attain high robustness with a corresponding trade-off in yield. And finally, the range of DNA concentrations that give rise to high yield and high robustness are often quite narrow indicating that feedback regulation may become a necessary design requirement [44].

## Discussion

We demonstrate how a biochemical constructor could be created by implementing a transcriptiontranslation system running at steady-state on a micro-chemostat that supplies the reaction with resources and energy. We showed that the system is capable of self-regenerating components of its core constructor by synthesizing proteins required for transcription and translation. We regenerated up to seven components simultaneously and show that system optimality is surprisingly similar to fitness landscapes observed in living systems [45], requires both minimizing resource loading and optimizing resource allocation, and can be described in terms of Pareto optimality.

Just like the universal constructor envisioned by von Neumann ~80 years ago, a biochemical universal constructor will consist of 3 components: i) an instruction set (DNA), ii) a core constructor (RNA and proteins), iii) and a copy machine (proteins). The core constructor consists of RNAs and proteins that read and implement the information contained in the instruction set. The core constructor is capable of constructing copies of itself and of the copy machine. The copy machine consists of the protein components necessary for DNA replication which copy the instruction set [14]. Similar to von Neumann’s universal constructor, the biochemical constructor requires supply of resources and energy, which is also a necessary requirement for all living systems.

Although we show that creation of a biochemical constructor is feasible, a number of considerable challenges remain. It will be critical to develop a transcription-translation system with a high enough synthesis rate to self-regenerate all of its components. The PURE system is currently orders of magnitude away from this target. We estimate that around 50% of all PURE proteins could be regenerated by the current PURE system, and that the total synthesis rate required is 25 fold above the current rate (Supplementary Fig. S22). These estimates do not yet include ribosome or tRNA synthesis. Current approaches to optimizing transcription-translation systems mainly focus on increasing component concentrations or adding components to the system which can give rise to overall higher synthesis yields but consequently also require higher synthesis rates to achieve self-regeneration [24, 25, 31]. Instead, optimizing protein synthesis rates and the ratio of protein synthesis rate to total amount of protein contained in the system will be important for development of a biochemical universal constructor. A second major challenge lies in achieving functional *in vitro* ribosome biogenesis [46, 21]. The most promising near-term goal will be demonstration of steady-state self-replication of DNA. Several promising advances have recently been demonstrated in this area [14, 15], although in *vitro* DNA replication efficiency likely needs to be improved in order to reach sustained steady-state DNA replication.

Achieving high yield and robustness will be as well important for the development of a universal biochemical constructor. These concepts are tightly connected to resource usage and loading effects recently described in cell-free systems [38] and living cells [47]. We showed that several components could be regenerated at the same time. However, finding optimal DNA concentrations for several components is critical to achieving sustained regeneration without unnecessarily loading the system. Moreover, our results and corresponding modeling suggest that specific components might have to be tightly regulated, and could benefit from active feedback regulation [44], especially once system complexity increases. Currently, self-regenerating systems can be optimized by varying individual DNA input concentrations in order to adjust protein synthesis rates for each component being regenerated. In the future, all genes will be encoded on a single ‘genome’ [15, 48], requiring expression strengths to be tuned by the use of synthetic transcription factors [49], promoters [50], terminators [51], and ribosome binding sites [52]. Work on a biochemical universal constructor thus provides ample challenges and opportunities for synthetic biology in the areas of protein biochemistry, tRNA synthesis, ribosome biogenesis, metabolism, regulatory systems, genome design, and system engineering.

The development of a universal biochemical constructor and the creation of synthetic life are exciting prospects and recent progress in technology and biochemistry are making these seemingly plausible goals. Many challenges remain, but pieces to the puzzle are being added at an increasing rate. It is thus not far-fetched to consider that synthetic life, engineered by humans from basic building blocks, may be a possibility.

## Acknowledgments

This work was supported by Human Frontier Science Program Grant RGP0032/2015; the European Research Council under the European Union’s Horizon 2020 research and innovation program Grant 723106; and EPFL. N.L. is supported by a Chancellor’s Fellowship from the University of Edinburgh.

## Author contributions

B.L. performed experiments. N.L. performed the computational analysis. B.L., N.L., and S.J.M. designed experiments, analyzed data, and wrote the manuscript.

## Data deposition

All supporting data and code are available on GitHub at https://github.com/lbnc-epfl. Microfluidic design files are available at http://lbnc.epfl.ch.

## Supporting Information

### Materials and methods

#### Materials

*E. coli* BL21(DE3) and M15 strains were used for protein expression. *E. coli* RB1 strain [1] originally obtained from G. Church (Wyss Institute, Harvard University, USA) was used for His-tag ribosome purification. All plasmids encoding PURE proteins used in this work were originally obtained from Y. Shimizu (RIKEN Quantitative Biology Center, Japan). Plasmid encoding mScarlet was a gift from P. Freemont (Imperial College London, UK).

Linear template DNA for *in vitro* eGFP synthesis (Supplementary Table S6) was initially prepared by extension PCR from a pKT127 plasmid as described previously [2] and cloned into a pSBlue-1 plasmid. The DNA fragment used for PURE system characterization and self-replication experiments was amplified from this plasmid by PCR. Linear DNA fragments encoding different proteins used for self-regeneration experiments were prepared by extension PCR from their respective plasmids. Primer sequences are listed in Supplementary Table S7. All DNA fragments were purified using DNA Clean and Concentrator-25 (Zymo Research). DNA was eluted in nuclease-free water instead of elution buffer, and its concentration was quantified by absorbance (NanoDrop, ThermoFisher). Double stranded Chi DNA [3] was prepared by annealing to primers listed in Supplementary Table S6.

#### Ribosome purification

All buffers used in this work are listed in Supplementary Table S8. All buffers were filtered (Flow Bottle Top Filters, 0.45 μm aPES membrane) and stored at 4°C. 2-mercaptoethanol was added immediately before use. Ribosomes were prepared from *E. coli* RB1 strain by His-tag purification [1]. *E. coli* RB1 strain was grown overnight in 3 mL LB media at 37°C. 4 × 3 mL of the overnight culture was used to inoculate 4 × 500 mL of LB in a 1 L baffled flask. Cells were grown at 37°C, 260 RPM to exponential phase (3-4 h), pooled together and harvested by centrifugation (4,000 rpm, 20 min, at 4°C), and stored at −80°C. The cells were then resuspended in 15 mL suspension buffer and lysed by sonication on ice (Vibra cell 75186, probe tip diameter: 6 mm, 11 × 20s:20s pulse, 70% amplitude). Cell debris was removed by centrifugation (15000 rpm, 20 min, at 4°C). The recovered fraction was filtered with a GD/X syringe filter membrane (0.45 mm, PVDF, Whatman).

Ribosomes were purified using 5 mL IMAC Sepharose 6 FF (GE Healthcare) by Ni-NTA gravity-flow chromatography. The corresponding buffers were prepared by mixing buffer C and buffer D at the required ratios. After the column was equilibrated with 30 mL of lysis buffer (100% buffer C), the prepared lysate solution was loaded onto the column. The column was washed with 30 mL of lysis buffer (100% buffer C), followed by 30 mL of wash buffer 1 (5 mM imidazole), 60 mL of wash buffer 2 (25 mM imidazole), 30 mL wash buffer 3 (40 mM imidazole), 30 mL wash buffer 4 (60 mM imidazole) and eluted with 7.5 mL elution buffer (150 mM imidazole). Ribosomes from two purifications were pooled together (around 15 mL) and subjected to buffer exchange using a 15 mL Amicon Ultra filter unit with a 3 kDa molecular weight cutoff (Merck). All centrifugation steps were performed at 4000 rpm and 4°C. The elution fraction was concentrated to 1 mL (60 min). The concentrated sample was then diluted with 15 mL of ribosome buffer and re-concentrated to 1 mL (60-70 min); this step was repeated three times. The recovered ribosomes (1 mL) were further concentrated using a 0.5 mL Amicon Ultra filter unit with a 3 kDa molecular weight cutoff (Merck) by centrifugation (14,000 RCF, at 4°C). Ribosome concentration was determined by measuring absorbance at 260 nm of a 1:100 dilution. An absorbance of 10 for the diluted solution corresponds to a 23 *μM* concentration of undiluted ribosome solution. Final ribosome solution used for *in vitro* protein synthesis was prepared by diluting to 3.45 *μ*M. The usual yield is around 0.75 mL of 3.45 *μ*M ribosome solution.

#### Ni-NTA resin preparation and regeneration for ribosome purification

5 mL IMAC Sepharose 6 FF (GE Healthcare) was pipetted into Econo-Pac chromatography columns (Bio-Rad), and charged with 15 mL of 100 mM nickel sulfate solution. The charged column was washed with 50 mL of demineralized water. After protein purification, columns were regenerated with 10 mL of buffer containing 0.2 M EDTA and 0.5 M NaCl, and washed with 30 mL of 0.5 M NaCl, followed by 30 mL of demineralized water, and stored in 20% ethanol at 4°C.

#### PURE system preparation

Proteins were purified by Ni-NTA gravity-flow chromatography as described previously [4]. Different PURE protein formulations are summarised in Supplementary Table S3. Different PURE or ΔPURE systems were prepared by supplying the corresponding ΔPURE systems with the omitted protein or buffer solution, respectively.

#### Energy solution preparation

Energy solution was prepared as described previously with slight modifications [4]. 2.5× energy solution contained 0.75 mM of each amino acid, 29.5 mM magnesium acetate, 250 mM potassium glutamate, 5 mM ATP and GTP, 2.5 mM CTP, UTP and TCEP (tris(2-carboxyethyl)phosphine hydrochloride), 130 U_*A*260_/mL tRNA, 50 mM creatine phospate, 0.05 mM folinic acid, 5 mM spermidine, and 125 mM HEPES.

#### Batch *in vitro* protein expression experiments

Batch PURE reactions (5 *μ*L) were established by mixing 2 *μ*L of 2.5× energy solution, 0.9 *μ*L of 3.45 *μ*M ribosomes (final concentration: 0.6 *μ*M), 0.65 *μ*L of PURE proteins (Supplementary Table S3), DNA template, and brought to a final volume of 5 *μ*L with addition of water. All reactions measuring eGFP expression were prepared as described above with eGFP linear template at a final concentration of 4 nM and incubated at 37°C at constant shaking for 3 h, and measured (excitation: 488 nm, emission: 507 nm) on a SynergyMX platereader (BioTek). The eGFP production rate was calculated between 20-50 min based on an eGFP calibration curve (Supplementary Fig. S23A). Reactions expressing other proteins were prepared as described above and supplemented with 0.2 *μ*L FluoroTect GreenLys (Promega). DNA templates were used at a final concentration of 2 nM and the reactions were incubated at 37°C for 3 h.

#### SDS-PAGE gels

PURE reactions (5 *μ*L) labeled with FluoroTect GreenLys (Promega) were incubated with 0.8 μg or 0.2 *μ*L of RNAse A solution (Promega) and incubated for 30 min at 37°C and subsequently analyzed by SDS-PAGE using 10-well 4-20% Mini-PROTEAN TGX Precast Protein Gels (Bio-Rad). Gels were scanned (AlexaFluor 488 settings, excitation: Spectra blue 470nm, emission: F-535 Y2 filter) with a Fusion FX7 Imaging System (Vilber) and analyzed with ImageJ. Protein sizes were calculated based on a BenchMark™ Fluorescent Protein Standard (Invitrogen).

#### Fabrication and design of the microfluidic device

The microfluidic device was fabricated by standard multilayer soft lithography [5], detailed device preparation, operation, and characterisation are described previously [6]. The device with 8 reactors and 9 fluid inputs (Fig. S1) is based on a previous design [2].

#### Device setup

To prime the chip, control lines were filled with phosphate buffered saline (PBS) and pressurized at 1.38 bar. The flow layer was primed with a solution of 2% bovine serum albumin (BSA) in 0.5× PBS. For washes between loading steps, 10 mM TRIS buffer (pH = 8) was used. For the experiments energy, PURE, and DNA solutions were mixed in the microfluidic reactors on the microfluidic chip in a 2:2:1 ratio. The peristaltic pump was actuated at 20 Hz to mix the solutions. Every 15 min, the reactor was imaged and a 20% fraction of the reactor volume was replaced with fresh components with the same 2:2:1 ratio. Details on the operation of the microfluidic chip can be found in Supplementary Tables S1 and S2. 2.5× energy solution was prepared as described above. 2.5× PURE or ΔPURE solutions were prepared by mixing the desired protein solutions (Supplementary Table S3) with ribosomes (final concentration: 0.6 *μ*M) and supplied with 10 *μ*M TCEP (final concentration: 4 *μ*M). The PURE solution was supplemented with mScarlet protein to allow visualization, and the solutions were brought to final volume with the addition of water. The DNA solution at five times its final concentration was prepared by mixing the desired linear templates and Chi DNA. The final concentration of eGFP reporter in the reaction was 2 nM, the Chi DNA was used at a final concentration 1.25 *μ*M.

#### Data acquisition and analysis

Solenoid valves, microscope, and camera were controlled by a custom LabVIEW program. The chip and microscope stage were enclosed in an environmental chamber at 34°C. Green and red fluorescence was monitored over time on an automated inverted fluorescence microscope (Nikon), using 20x magnification and FITC / mCherry filters. The microscope hardware details are described in [6].

The fluorescence images were analyzed and corrected in Python, by subtracting the background fluorescence of a position next to the fluidic channel. The fluorescence signal was normalized in respect to maximal positive control signal intensity in a given experiment, or to the overall maximal intensity if a positive control was not included. The eGFP synthesis rate was calculated based on an eGFP calibration curve (Supplementary Fig. S23B) and dilution rate.

### Modeling

#### Minimal resource-dependent TX-TL model

While cell-free transcription and translation can be described at varying levels of granularity [7, 8, 9, 10, 11, 12, 13, 14], here we chose to model the processes at the most coarse-grained level using coupled ordinary differential equations (ODEs). This model can be easily extended to incorporate more complex effects, but the aim here was to show a minimal mechanism which qualitatively captures the observed experimental effects.

The model consists of simultaneous transcription and translation of GFP and T7 RNAP, which consumes a single resource species *R*. This species is a lumped representation of NTPs which are consumed during transcription, and ATP, GTP, and aminoacyl tRNAs which are consumed during translation. We model the transcription rate by a parameter *α*, linearly dependent on DNA and T7 RNAP concentration, and modulated by the availability of resources using a Hill function *R*/(*R* + *K*). Likewise, translation proceeds at a rate *β*, which is linearly dependent on mRNA concentration, and is modulated by the same Hill function for resource dependence. The rate of consumption of R is equal to the summed transcription and translation rates. The complete model consisting of seven ODEs and three parameters, is shown below.

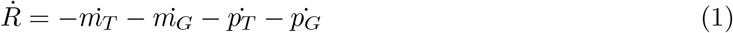

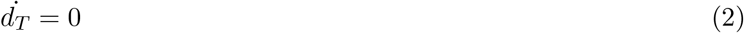

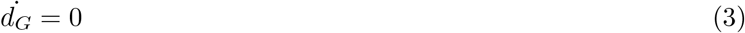

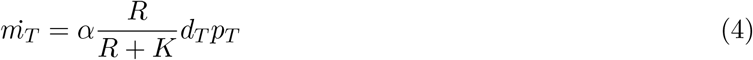

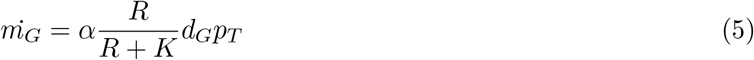

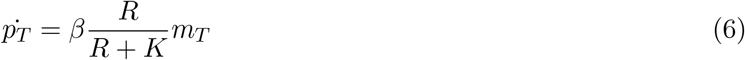

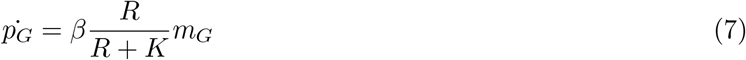

DNA, mRNA, and protein concentrations are denoted by *d*, *m*, and *p*, and the subscripts *T* and *G* refer to T7 RNAP and eGFP respectively. The model was implemented in Julia 1.4.2 and solved using the `DifferentialEquations.jl` package. All code is available on github.

#### Chemostat simulation

During chemostat operation, concentrations of species in the cell-free reaction are periodically adjusted. All components are diluted at a specific dilution fraction, while certain components (proteins, ribosomes, energy solution, and DNA) are replenished. This can be captured in the model by explicitly including the dilution steps. In Julia this is achieved by implementing callbacks which modify the concentrations at specified time points while solving the ODEs. More detail can be found in the documentation for the code.

Simulating the chemostat leads to a sawtooth-like behaviour as eGFP is diluted, and subsequently produced between dilution steps, as shown by the green curve in Supplementary Figure S9. In the real experiment, images are taken immediately before each dilution step, and thus the data appear smooth: this is shown by the dashed black curve in Supplementary Figure S9.

At each dilution step in the model, all species’ concentrations are reduced by a fixed dilution fraction, while the concentrations of certain species are refreshed by addition of a fraction of those species at their initial concentrations:

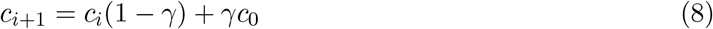

The dilution fraction *γ* was set to 20%, and the periodicity of dilution to 15 minutes, corresponding to experimental values. Each *in silico* experiment contained three stages: 1.) kick-start, 2.) self-regeneration, and 3.) washout. The species were replenished as indicated in Supplementary Table S5. The negative control corresponded to a self-regeneration experiment with *d_T_* = 0.

#### Model design and parameters

Since our overall goal was to capture qualitative rather than quantitative behaviour, we used effective parameters and arbitrary units to describe system dynamics, which were combined with physical time values and experimental chemostat operation parameters. Nevertheless, initial parameter selection was guided by relative magnitudes of various parameters. In particular, the resource saturation term *K* is typically several orders of magnitude less than the initial resource concentration [15], and cell-free translation rates are typically an order of magnitude slower than transcription [16]. The initial parameter set was manually chosen to reflect experimentally-observed behaviour; parameter scans were then conducted to test the robustness of model behaviour on parameter variations, as discussed below.

We have assumed equal resource consumption for transcription and translation, which is motivated by the fact that translation consumes 2*N* GTP and *N* ATP to synthesise a polypeptide of length N (accounting for aminoacylation), while transcription consumes on average 3*N*/4 ATP and 3*N*/4 GTP (as well as UTP and CTP), which is the same as translation to within an order of magnitude.

#### Elucidation of model behaviour

In order to elucidate the origins of the observed behaviour, we can inspect protein, mRNA, and resource levels, as shown in Supplementary Figure S10. Because translation rate is the product of mRNA concentration and the resource dependence term *R*/(*R* + *K*), high levels of translation require both to be present. Let us consider the self-regeneration phase (4–16h). At low concentrations of *d_T_*, resource levels are high; however mRNA concentrations are low, and thus overall the translation rate of eGFP is low. In the converse situation at high *d_T_*, mRNA levels are high, but resources are low, leading once again to low eGFP production. It is only at intermediate concentrations of *d_T_* where eGFP production is high when there is a small but nonzero amount of resource availability, as well as an intermediate level of mRNA present. The model thus predicts that the production of eGFP is determined by a trade-off between resource availability and mRNA concentration.

In order to further interrogate the model, we can look at transcription and translation rates. These can be determined from the model by evaluating the derivatives directly from the ODEs. As the system is periodically diluted, the rates are also periodically modulated. An example is shown in Supplementary Figure S11A (grey line), from which we can calculate the average rate (green line). Here we observe that the translation rate of GFP in the positive control experiment varies from a high value to zero in every cycle. This is due to resources being completely depleted in each cycle, as shown in Supplementary Figure S11B. The average rates of transcription and translation of GFP and T7 RNAP are shown in Supplementary Figure S12, where we again observe that increasing *d_T_* increases average transcription rates (Fig. S12B), but decreases translation rates (Fig. S12A).

In our model the consumption of resources is directly equal to the summed transcription and translation rates. Thus we can determine the allocation of resources between different model processes, as shown in Supplementary Figure S13. We observe that at the onset of the self-regeneration phase, transcriptional resource consumption decreases while translational consumption increases (Fig S13A and B), which forms one part of our hypothesis to explain the increase in eGFP production over the positive control. Fig S13E and F show that as T7 RNAP is washed out at late times, resource allocation tends to 100% translation, and this accounts for the ‘bump’ in eGFP production in the washout phase.

We can also carry out parameter variations, shown in Supplementary Figure S14 as a contour map of the variation of the parameter of interest against a T7 DNA titration. Here we show the model predictions for two quantities: the ratio of self-regeneration to positive control (SR/PC) at 15h, as shown in the main text, and the overall eGFP production at 15h, which is a measure of the productivity of the self-regeneration process. These results yield further insights into the mechanisms of the model; of relevance is the observation that both eGFP production and the ratio SR/PC exhibits an optimum with respect to T7 RNAP DNA, and the position of the optimum is only significantly affected by transcription and translation rates: increasing these rates shifts the optimum to lower values. The optimum is otherwise relatively robust; in particular, the position of the SR/PC optimum is insensitive to *d_G_* (shown in more detail in Supplementary Figure S15). An important observation is that the the ratio SR/PC contains an optimum for high values of T7 DNA and small values of the initial resource concentration *R*_0_. The interpretation of this is that the model predicts that high SR/PC ratios are achieved when the resources become scarce.

In summary, analysis of the single resource-dependent model behaviour leads to two main conclusions:

1. eGFP production depends on a trade-off between resource availability and mRNA concentration. As *d_T_* is increased, eGFP production therefore exhibits a maximum.
2. For intermediate concentrations, eGFP production is higher than the positive control during the self-regeneration phase. This is accounted for by a reallocation of resources from transcription to translation during the transition between kick-start and self-regeneration, and by an overall resource-limited condition.

#### Sufficiency of model mechanism

We would like to understand whether the resource-dependent model is necessary and sufficient to explain our observations. Therefore we developed a second model, whose transcriptional and translational activities do not depend on any resource. This ‘resource-independent’ model instead contains TX and TL rates which decrease exponentially over time, with a fixed decay constant λ, which represents a resource-independent inactivation of cell-free protein synthesis. Such effects are also observed experimentally [8]. This model can be written as follows:

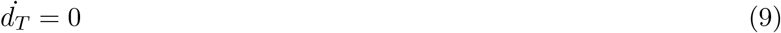

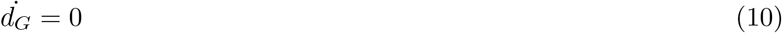

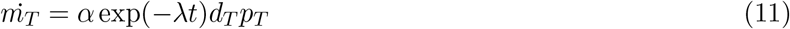

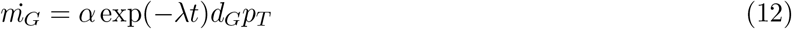

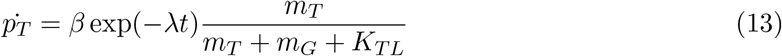

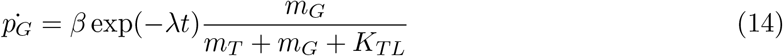

Here, we model translation as saturating at high total mRNA concentrations, with a Hill function and a saturation constant *K_TL_*. This is a typical way of taking into account translational loading effects [17]. We observe that this model can also qualitatively capture some of the experimental observations. Supplementary Figure S16A shows the time courses and SR/PC ratio plot of the single-resource model. Figure S16B shows the same for the resource-independent model, which again captures the optimum in SR/PC ratio as a function of T7 RNAP DNA. The model exhibits the three features of decaying eGFP production at low *d_T_*, high eGFP (potentially above the positive control level) at intermediate *d_T_*, and low eGFP production followed by a peak during washout at high *d_T_*.

The explanation of a maximum in eGFP production as a function of T7 RNAP DNA, is different from in the single-resource model. Here, at low *d_T_*, the concentration of *m_G_* is low, leading to low translation rates. At high *d_T_*, the concentration of *m_T_* is high, loading the translational machinery and again leading to low translation rates. At intermediate concentrations, where mRNA concentrations are high but before translational loading effects set in, we observe a maximum eGFP production.

Despite the different mechanism, there is a crucial similarity between this and the resource-dependent model: both involve coupling of the expression of eGFP and T7 RNAP. In the resource-dependent model, this is through a shared resource term, and in the resource-independent model, this is through a shared translational term. To demonstrate this, the coupling term can be artificially removed in the resource-independent model, allowing each protein to be translated independently. This leads to Figure S16C, where the ratio SR/PC monotonically increases with increasing T7 RNAP template, and no maximum is observed.

A second feature of both models is the striking increase of GFP above positive control levels (SR/PC> 1), for intermediate T7 RNAP template concentrations. These again result from two different mechanisms. In the resource-dependent model, analysis from the previous section shows that this is due to release of resources, under resource-limited conditions.

In the resource-independent model, T7 RNAP concentration is low in the kick-start phase. Thus any increase in T7 RNAP DNA template will increase T7 RNAP concentration, leading to greater transcription and translation, with no incurred costs. As long as translational capacity is not loaded, this effect can increase GFP over the positive control level. The increase of GFP is thus due to the activity of extra T7 RNAP in the system. However, the increase begins immediately in the kick-start phase, and is maintained throughout self-regeneration.

The explanation for these mechanisms can be tested *in silico*: in the first case, increasing the availability of resources should alleviate the resource constraint, and decrease the SR/PC ratio. This is observed in the parameter study shown in Figure S14. In the second case, increasing the initial T7 RNAP concentration should decrease the effect of any additional T7 RNAP produced. This is also observed in a similar parameter exploration, shown in Supplementary Figure S17.

In reality, it is likely that both mechanisms are at play. While the PURE system is known to be resource-limited under certain conditions [18], some lysate-based systems exhibit resource-independent deactivation [2]. Since a model which simultaneously takes both effects into account is likely to be more general, we tested a combined model, whose results are shown in Figure S16D and S18. While this model also successfully captures experimental observations, it is less robust than the simpler models, requiring fine-tuning of parameters. Experimentally, since we observe an increase in GFP after self-regeneration begins, and not immediately from the beginning of the kick-start phase, it is likely that under our experimental conditions, the resource limitation is a more dominant effect.

### Supporting Figures and Tables

**Figure S1:**
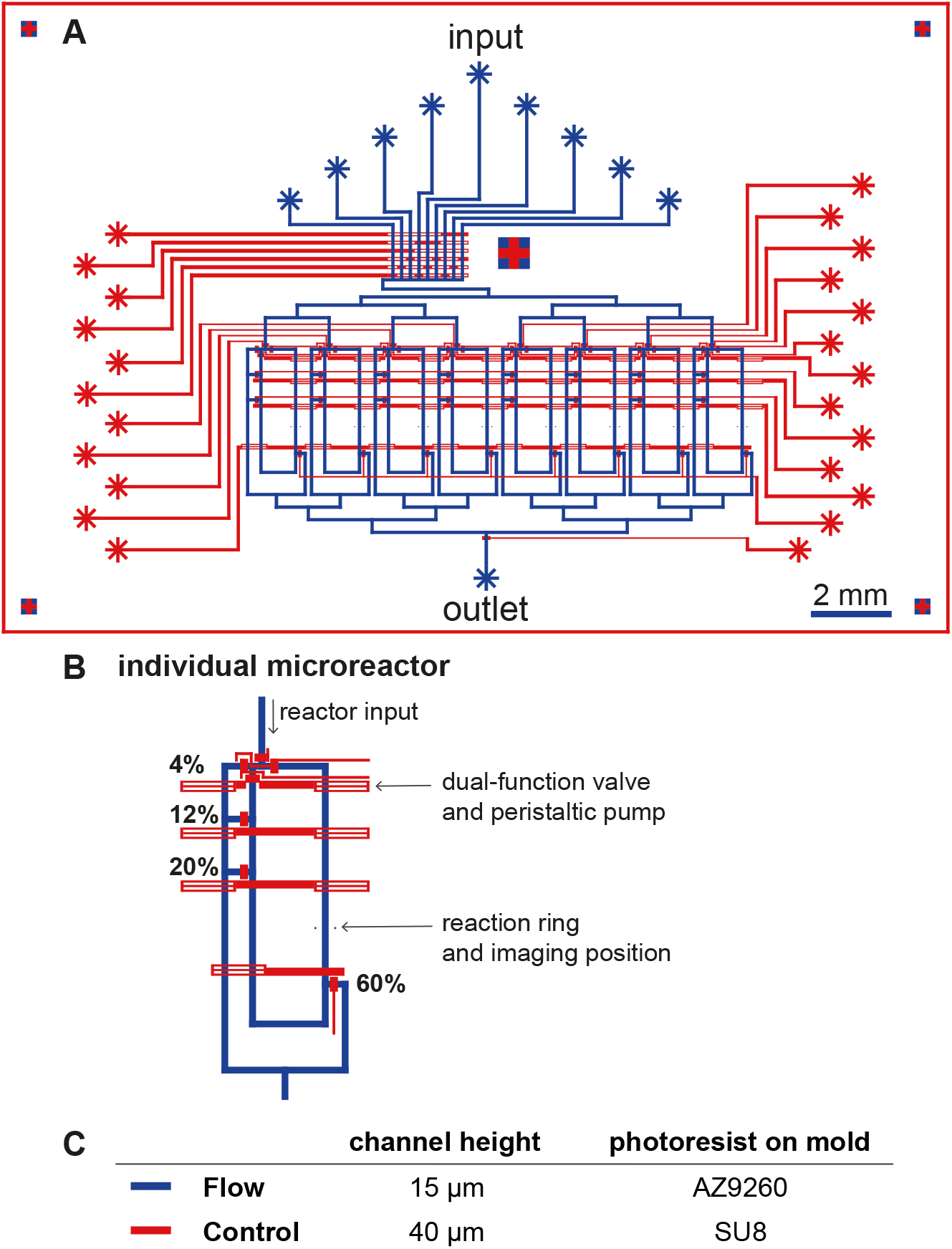
**(A)** Design schematic of the microfluidic device. The control layer is shown in red and the flow layer in blue. The device contains eight individually addressable chemostat reactors. **(B)** Close-up of a microfluidic reactor. Each reactor has four outlets corresponding to four different dilution fractions. Four control lines serve dual-functions as valves and peristaltic pump. The width of a flow channel is 100*μ*m. **(C)** Table of channel heights and corresponding photoresists used in mold fabrication.

**Figure S2:**
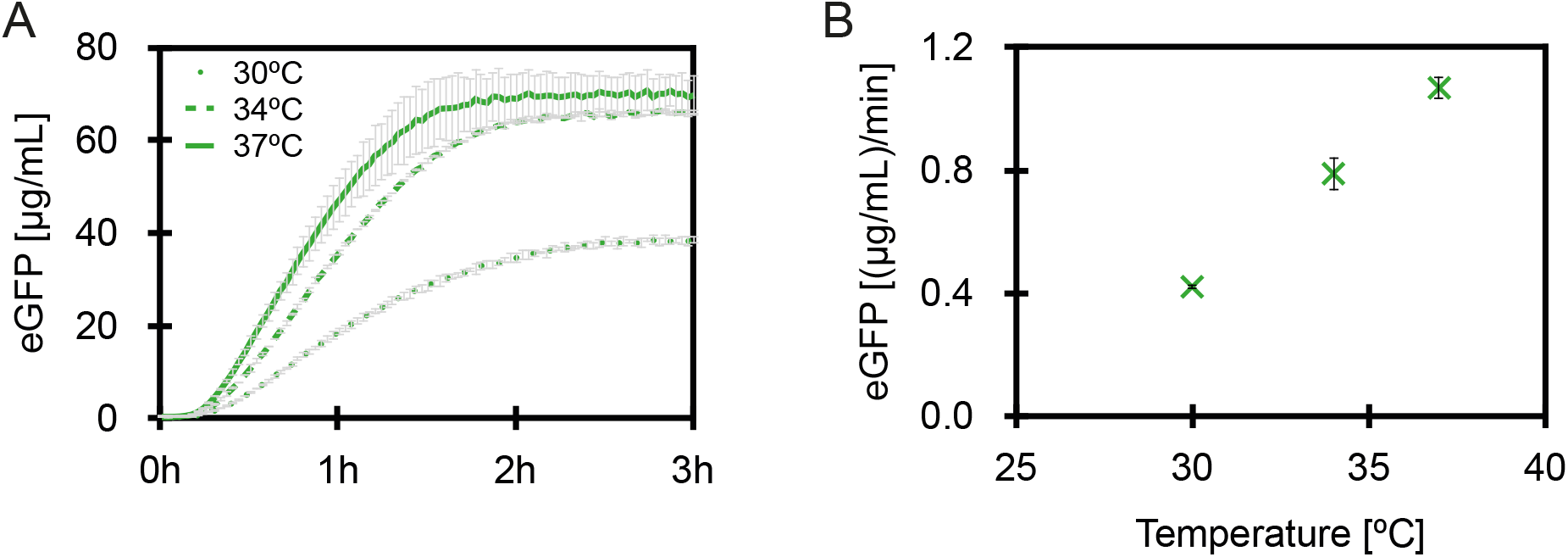
Comparison of eGFP expression at different temperatures in batch reactions, **(A)** eGFP expression over time, **(B)** eGFP expression rates. Each data point represents two technical replicates (mean ± s.d.).

**Figure S3:**
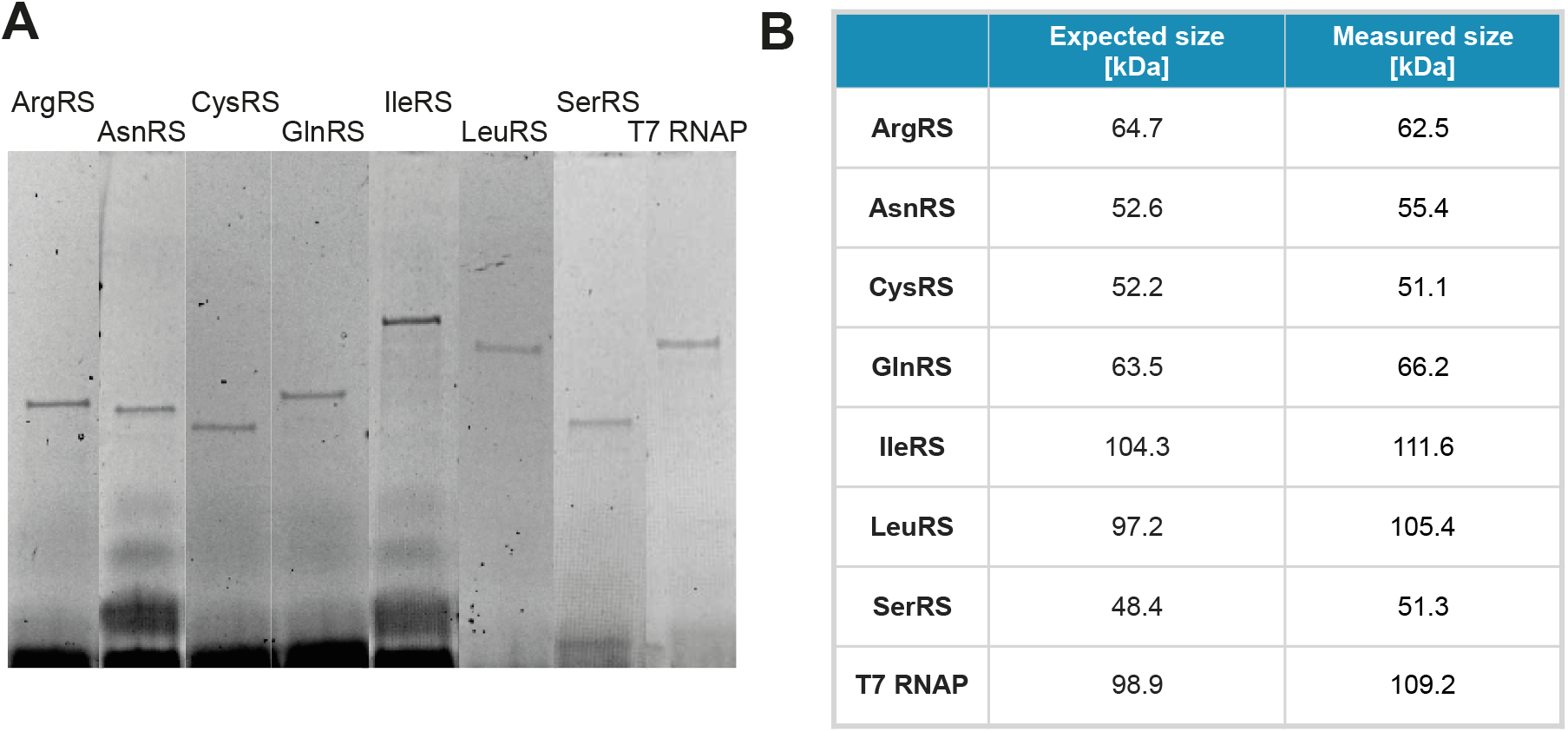
**(A)** SDS-PAGE gel of *in vitro* synthesized proteins labeled with FluoroTect GreenLys, **(B)** mass analysis of the expressed proteins.

**Figure S4:**
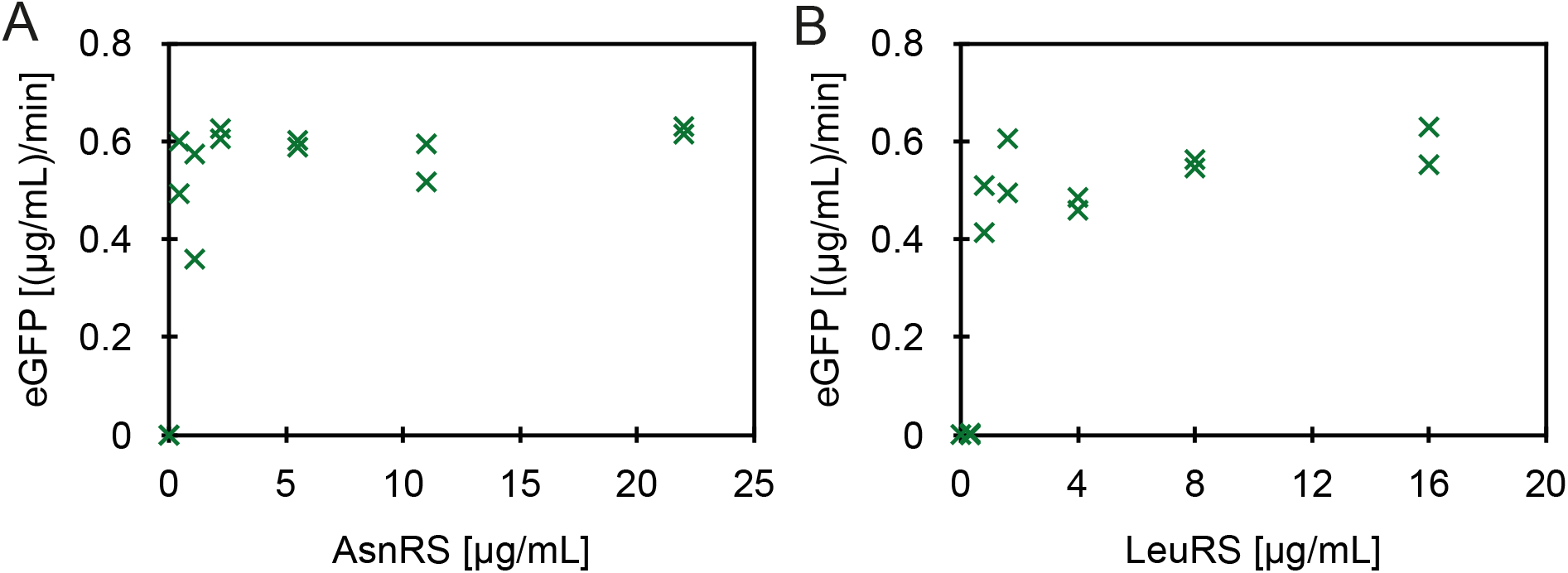
Comparison of eGFP expression rates in batch reactions at different components concentrations **(A)** AsnRS, **(B)** LeuRS. Each data point represents a technical replicate.

**Figure S5:**
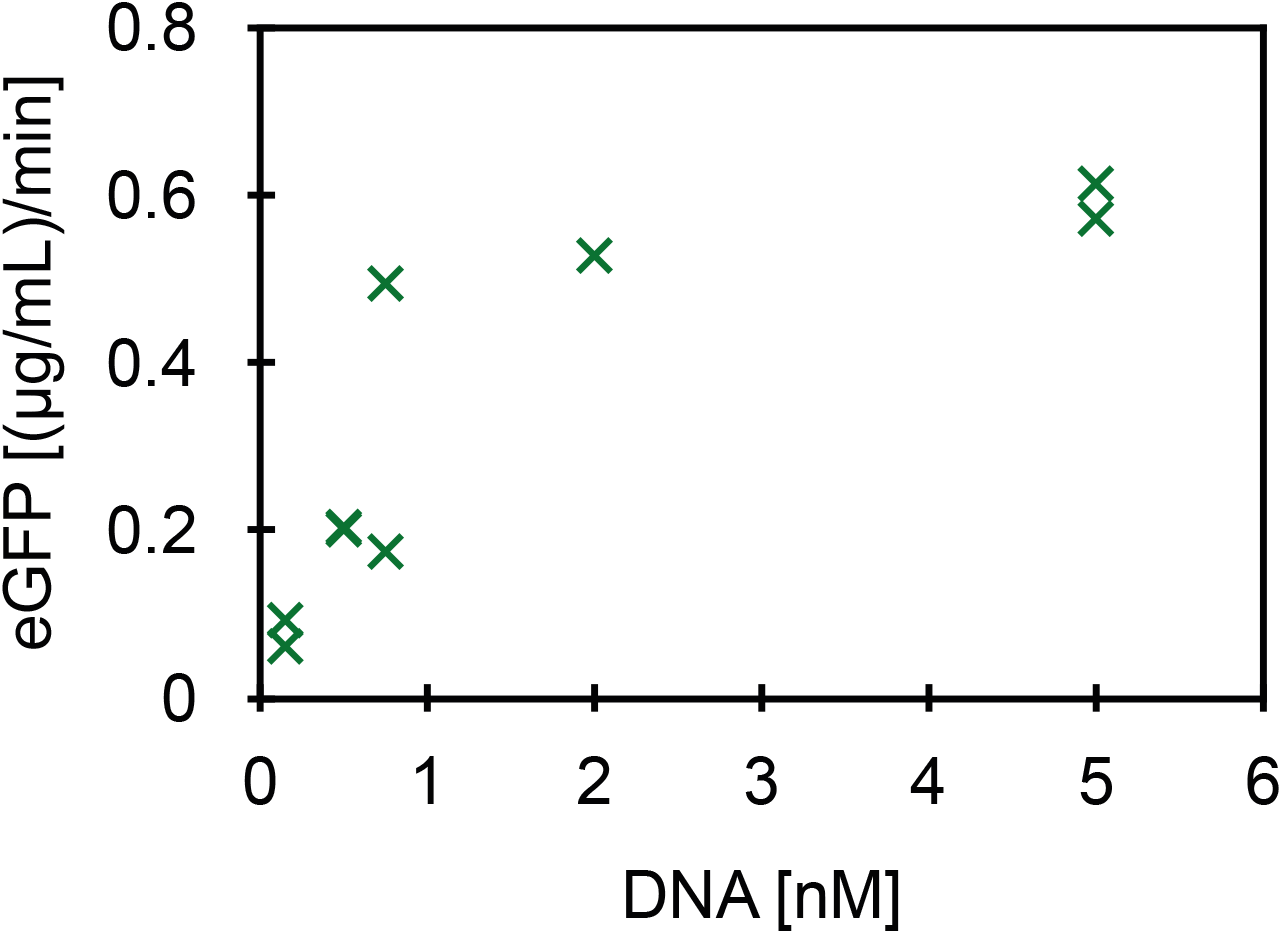
Comparison of eGFP expression rates in batch reactions at different DNA template concentrations. Each data point represents a technical replicate.

**Figure S6:**
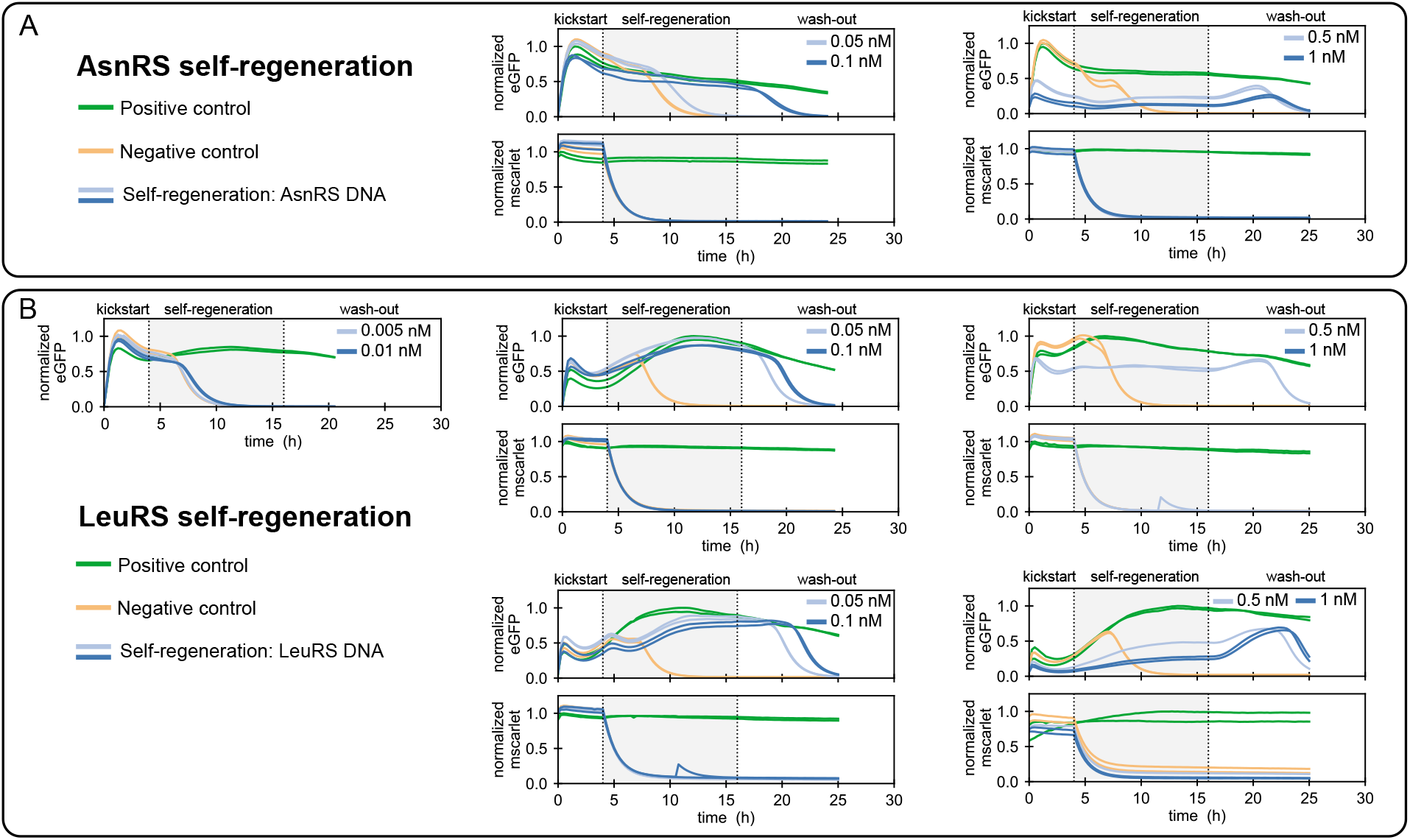
Summary of all **(A)** AsnRS and **(B)** LeuRS regeneration experiments and their corresponding mScarlet traces. The level of eGFP intensity is normalised to the maximum intensity obtained in the positive control. PURE system compositions used for the different experiments are given in Supplementary Table S3. 2 nM eGFP DNA template was used, and aaRS DNA template concentrations are indicated in the corresponding graphs.

**Figure S7:**
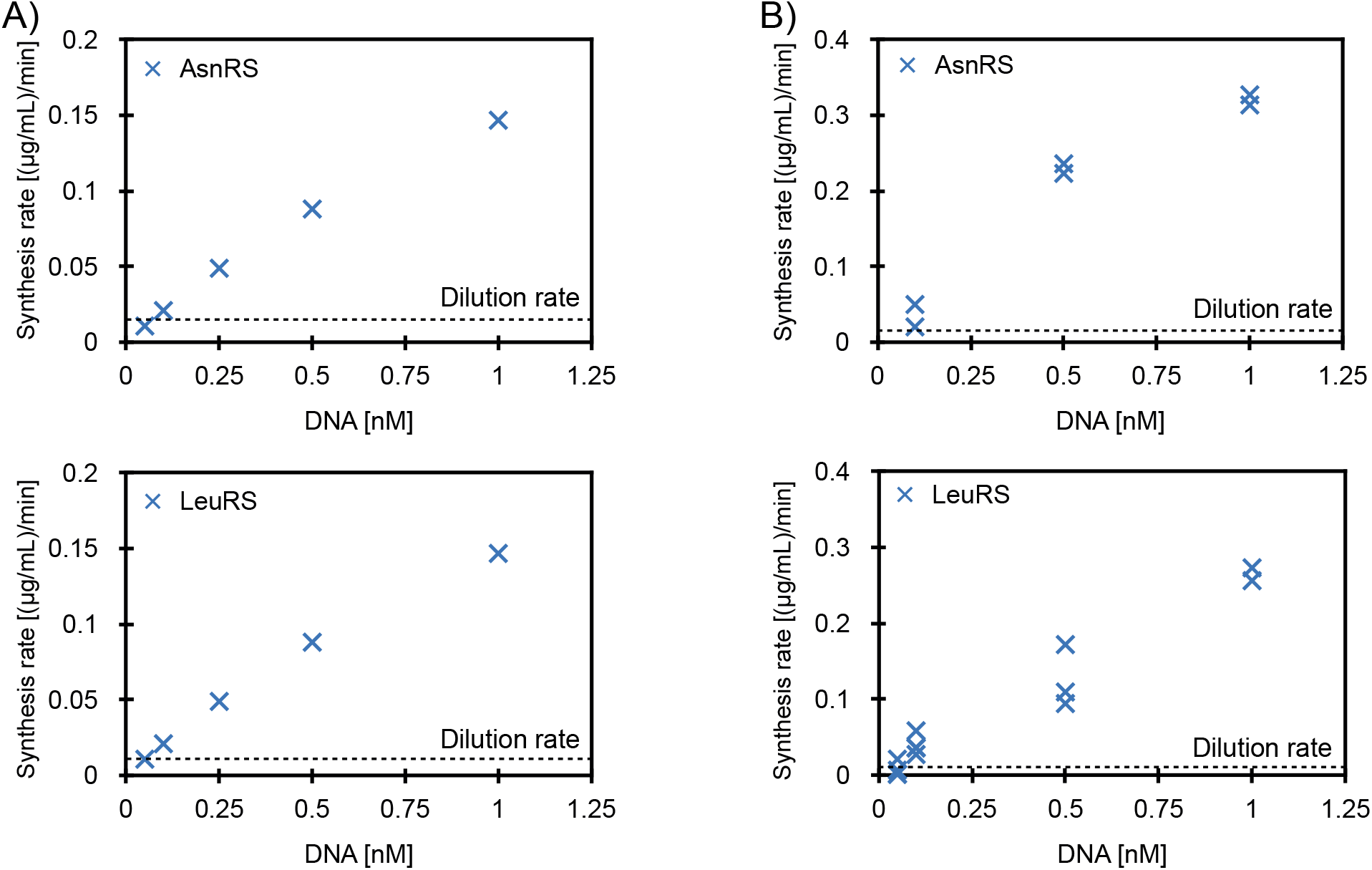
**(A)** Theoretical synthesis rate for single components expression, calculated based on eGFP synthesis rate in a microfluidic chemostat (0.44 (*μ*g/mL)/min) and DNA loading in DNA saturated system. **(B)** Estimated synthesis for AsnRS and LeuRS at different DNA concentrations based on the difference in eGFP synthesis rate for positive control and self-regeneration experiment at 15 hours. The eGFP synthesis rate was calculated based on an eGFP calibration curve (Supplementary Fig. S23B) and dilution rate. Dashed line represents the dilution rate of the given components based on the input component concentration (Supplementary Table S4).

**Figure S8:**
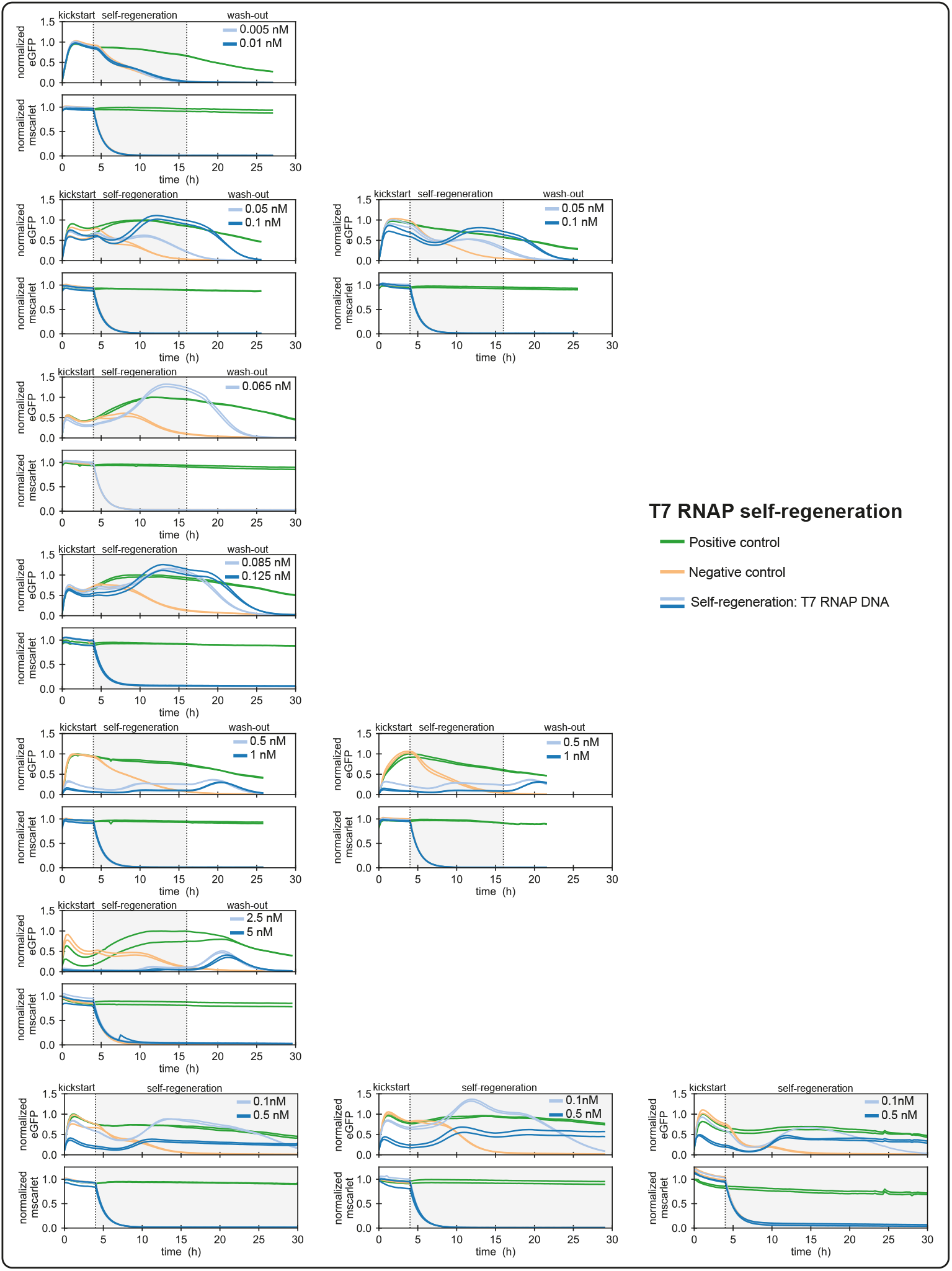
Results of regeneration experiments for all T7 RNAP DNA concentrations shown in Figure 2E, together with their corresponding mScarlet traces. The level of eGFP intensity is normalised to the maximum intensity obtained in the positive control. PURE system compositions used for different experiments are given in Supplementary Table S3. 2 nM of eGFP DNA template was used, and T7 RNAP DNA template concentrations are indicated in the corresponding graphs.

**Figure S9:**
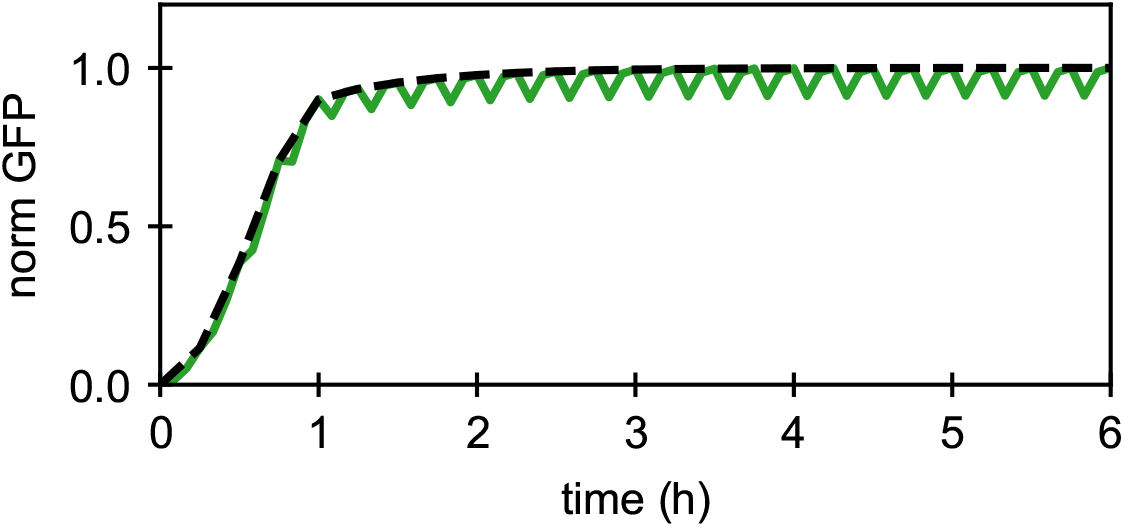
The chemostat is simulated by periodically diluting and replenishing species, and solving ODEs between the dilution steps. This leads to a sawtooth curve (green). Experimental measurements are taken immediately before each dilution step, which results in a smooth observation (dashed black line).

**Figure S10:**
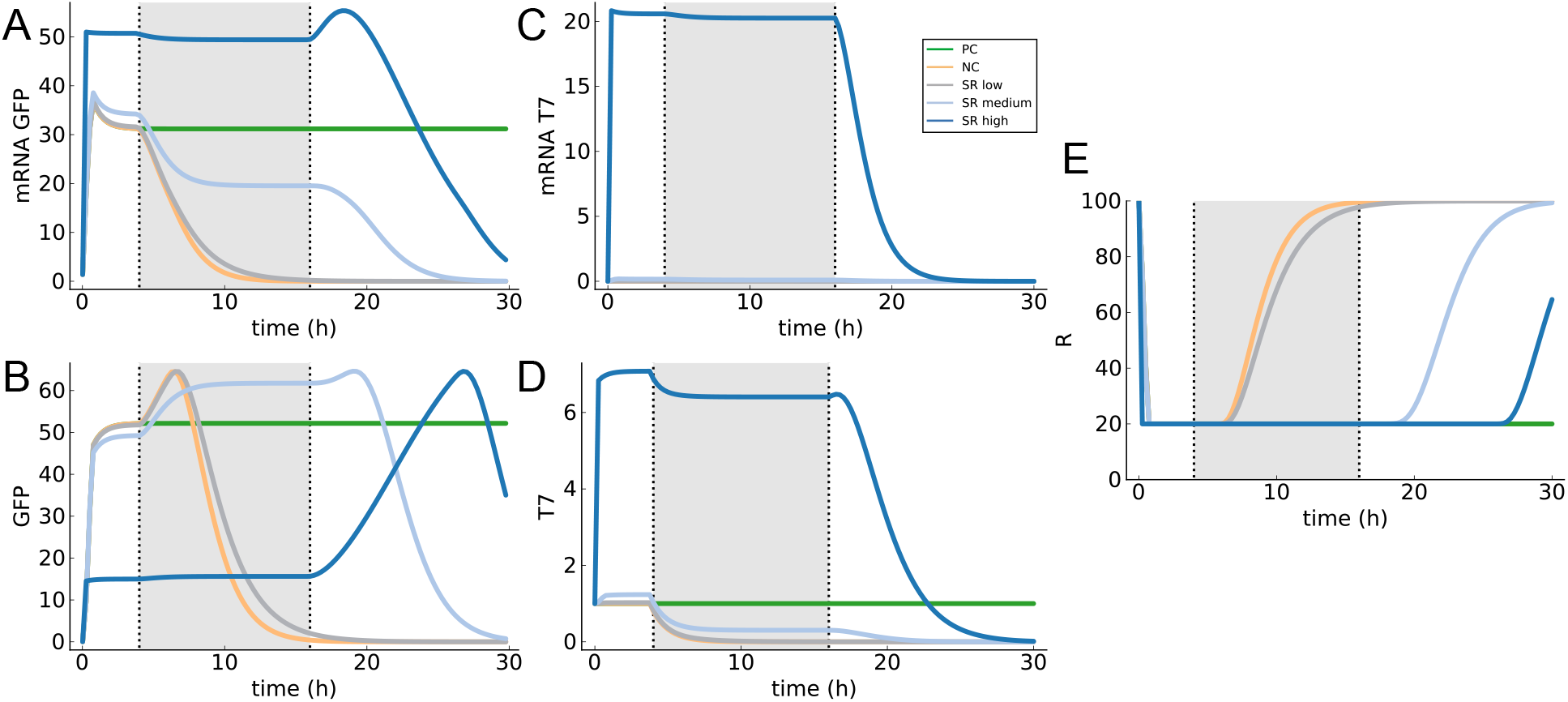
**(A,B)** Simulation results showing eGFP and **(C,D)** T7 RNAP mRNA and protein concentrations, as well as concentration of resource *R* **(E)**. Parameter values were *α* = 0.7, *β* = 0.07, *K* = 1 and initial conditions *R*_0_ = 100, *p_T_* = 1, *d_G_* = 2, with all other species set to zero. The three concentrations of *d_T_* are 0.001, 0.01, and 1, corresponding to the labels ‘low’, ‘medium’, and ‘high’, respectively.

**Figure S11:**
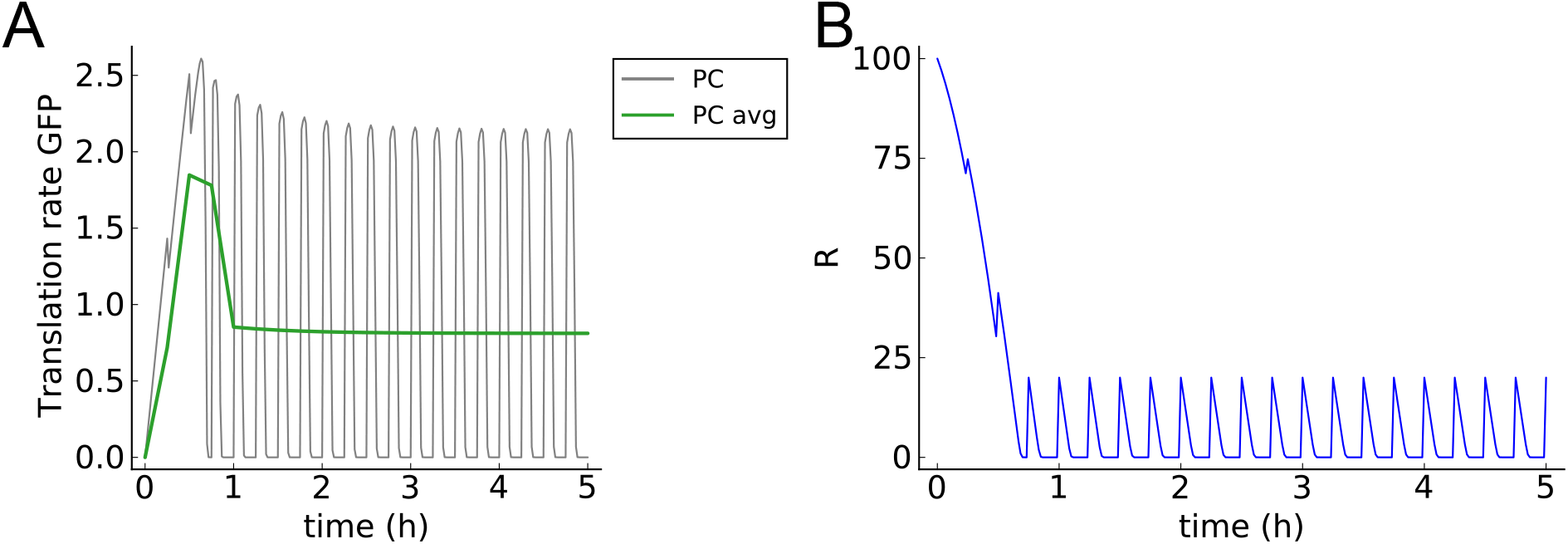
Derivatives can be directly calculated in the model, yielding rates of transcription and translation. **(A)** Periodic dilution of the chemostat leads to variations in rates, so we report the rates averaged over every period. **(B)** Translation of GFP occurs in a resource-limited regime, as resources are fully depleted over the course of each period.

**Figure S12:**
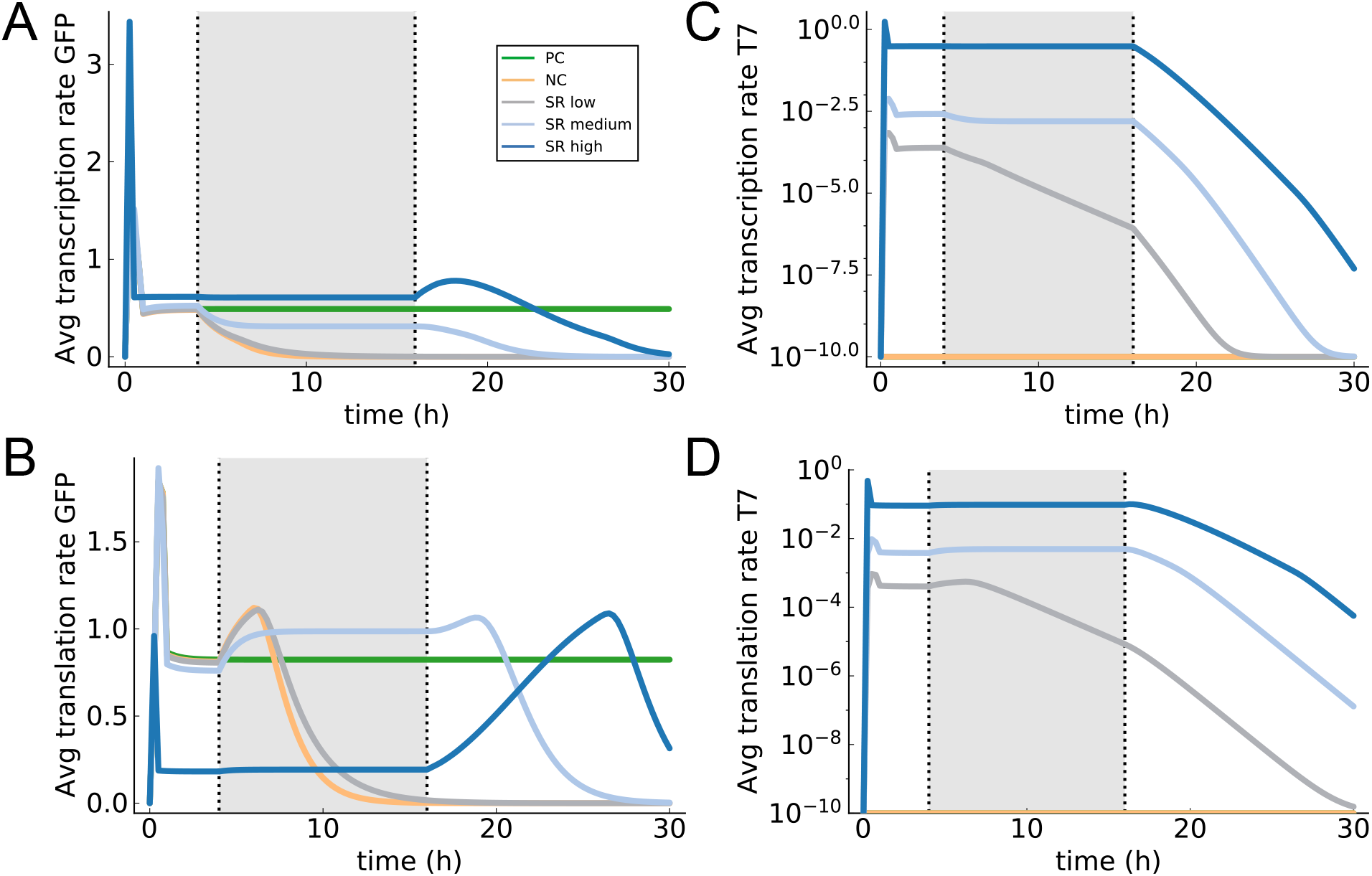
**(A,B)** Averaged transcription and translation rates for GFP and **(C,D)** T7 RNAP, for the same parameters as in Figure S10. To make the T7 rates more clear we plotted them on a log scale, with all values smaller than 10^−10^ set to 10^−10^.

**Figure S13:**
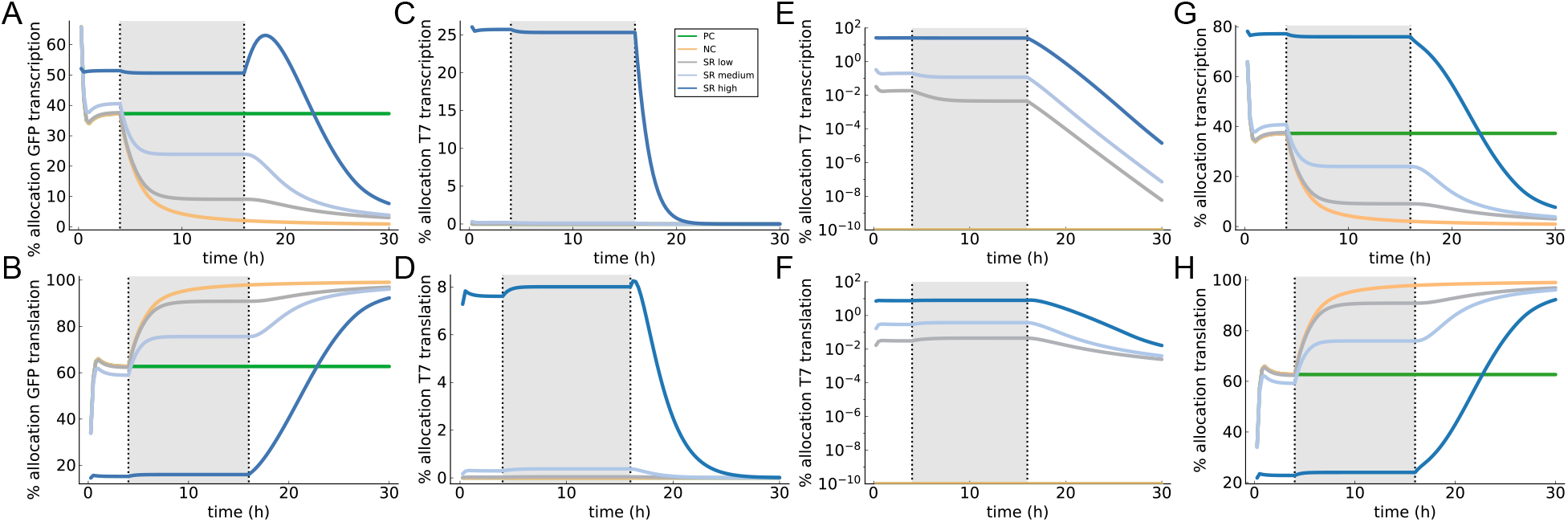
Varying resource allocation over the course of the simulation. **(A,B)** We observe reallocation of resources from transcription to translation at the beginning of the self-regeneration phase. **(C,D)** Resources consumed by T7 transcription and translation are shown on linear and **(E,F)** log scales for clarity. **(G,H)** The division of resources between total transcription and total translation. As T7 RNAP is washed out after 16 hours, resource allocation tends to 100% translation.

**Figure S14:**
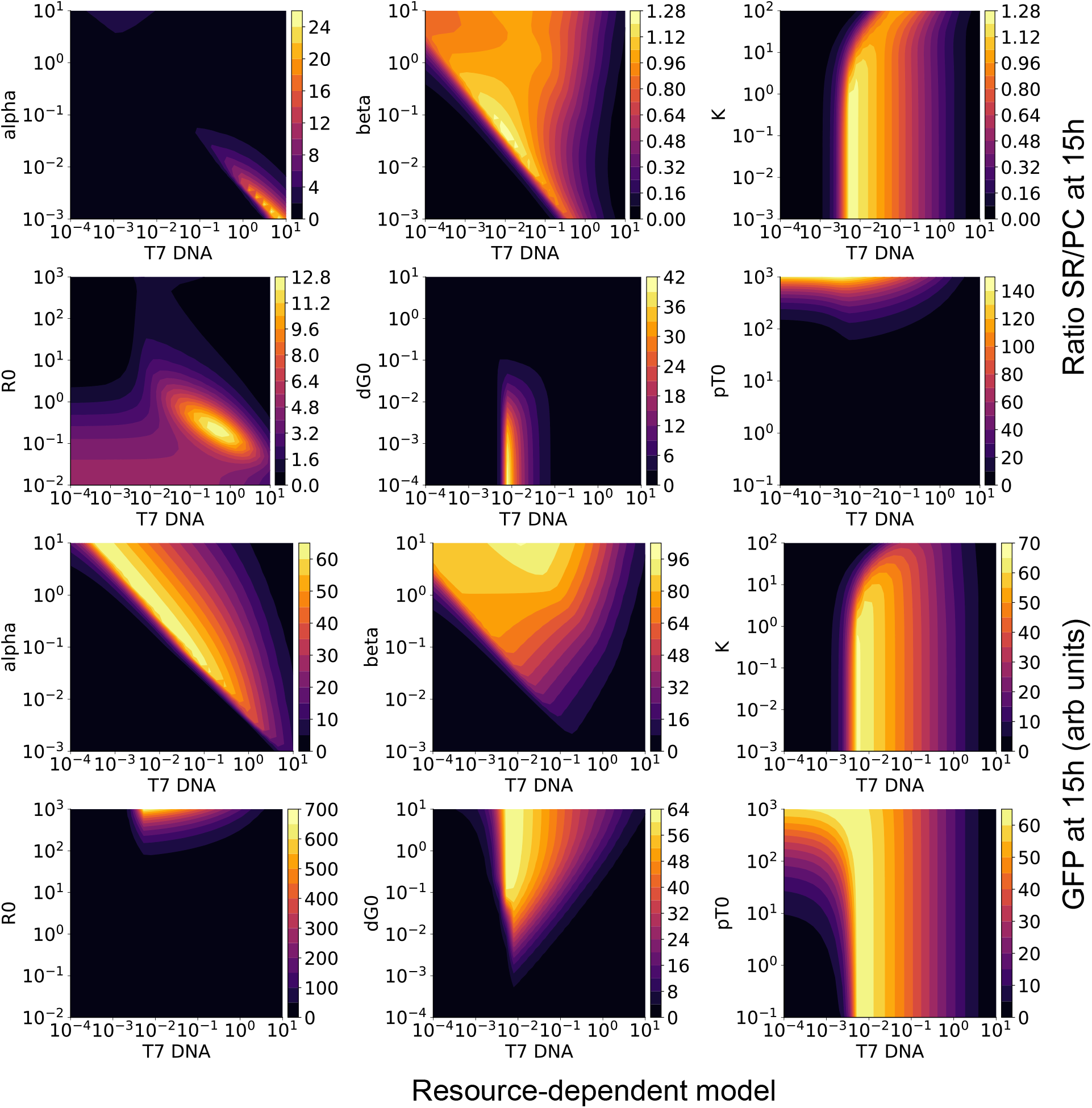
The effects of parameter variations on SR/PC ratio (top) and total GFP yield (bottom) for the resource-dependent model. We observe that SR/PC ratios are high for small values of *R*_0_, or for very high T7 DNA concentrations combined with low transcription rates; both these cases correspond to a resource-constrained situation.

**Figure S15:**
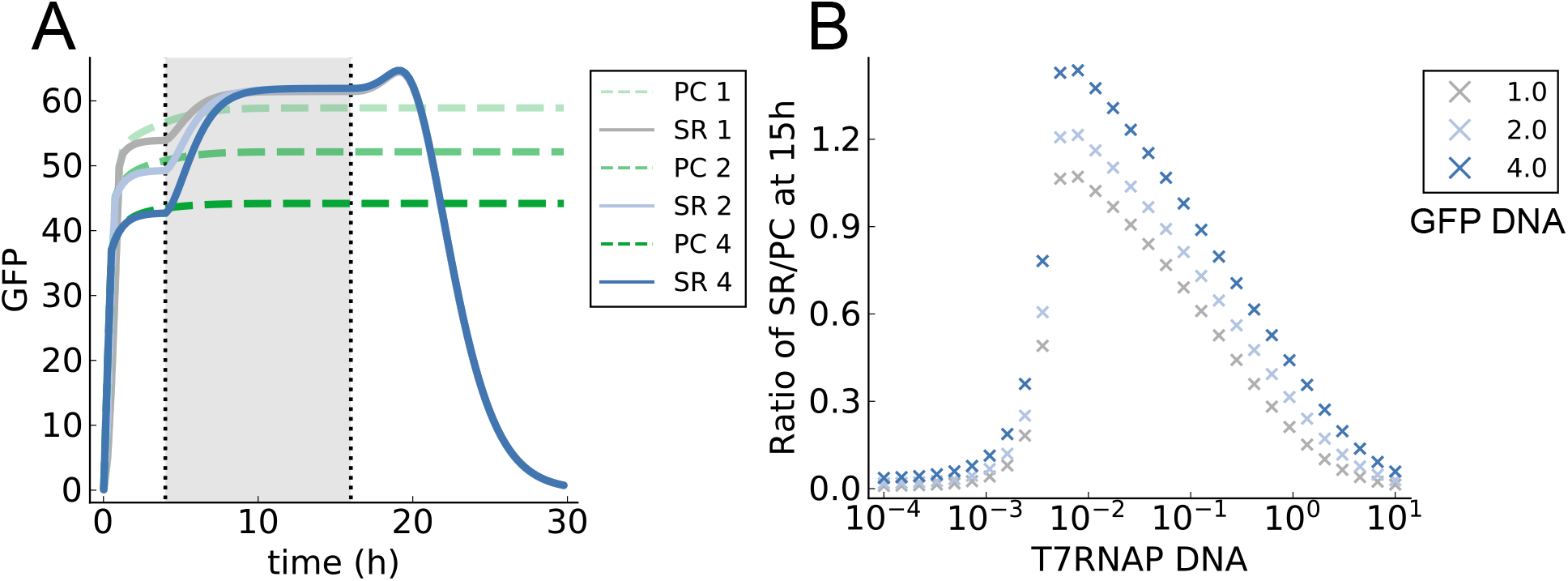
The effect of varying eGFP DNA (for values of 1, 2, and 4 nondimensional units) on the SR/PC ratio. **(A)** The resource-dependent model predicts that increasing eGFP DNA concentration lowers the positive control, as the reaction reaches steady state sooner due to faster consumption of resources. **(B)** This results in an increased SR/PC ratio during the self-regeneration phase.

**Figure S16:**
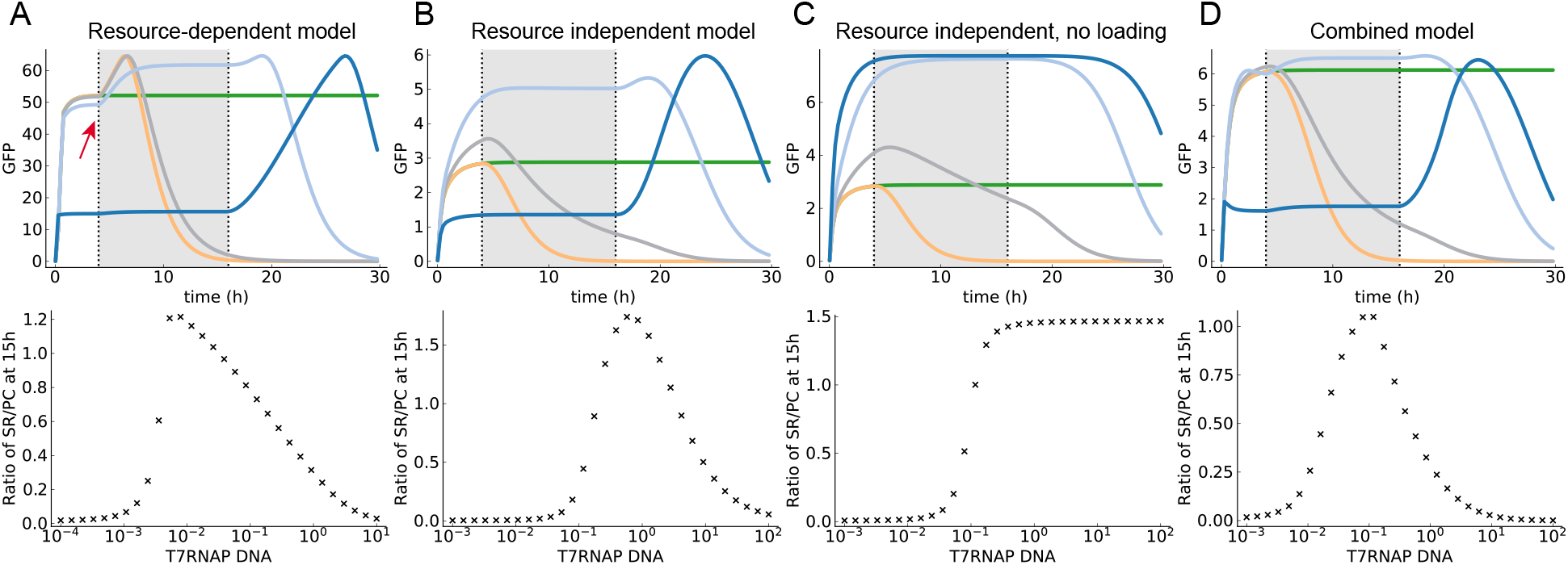
A resource-independent model **(B)** can also capture qualitatively similar results as the resource-dependent model **(A)**, showing a peak in eGFP production over a titration of T7 RNAP DNA. The major difference between the predicted behaviours is the rise in eGFP production after the beginning of the self-regeneration phase for the resource-dependent model (indicated by the red arrow), compared with the immediate rise at the beginning of the kick-start for the resource-independent model. Both models rely on coupling of eGFP and T7 RNAP production, through either a shared resource or enzyme. Removing the coupling eliminates the experimentally-observed optimum **(C)**. In reality both effects are likely to be present, and a combined model **(D)** can also capture experimental results, at the expense of increased complexity.

**Figure S17:**
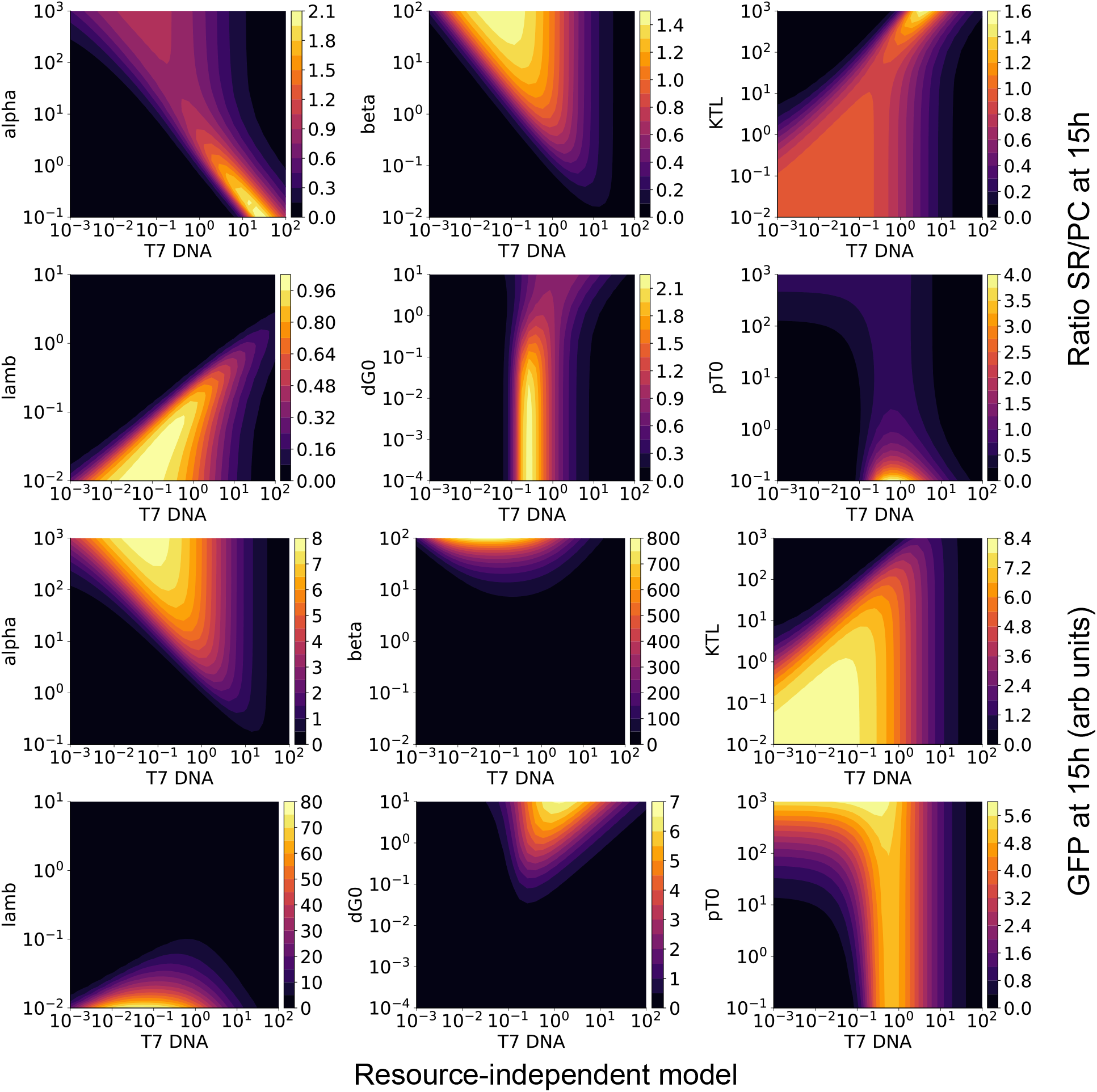
The effects of parameter variations on SR/PC ratio (top) and total GFP yield (bottom) for the resource-independent model. The variations are broadly similar to the resource-dependent model for the shared parameters *α, β, dG*_0_, and *pT*_0_. The behaviour of λ, the activity decay constant, is opposite to that of *R*_0_ for the single resource model, as both parameters qualitatively limit the reaction lifetime. Finally, the model is sensitive to variations in the translation saturation constant *K_TL_*.

**Figure S18:**
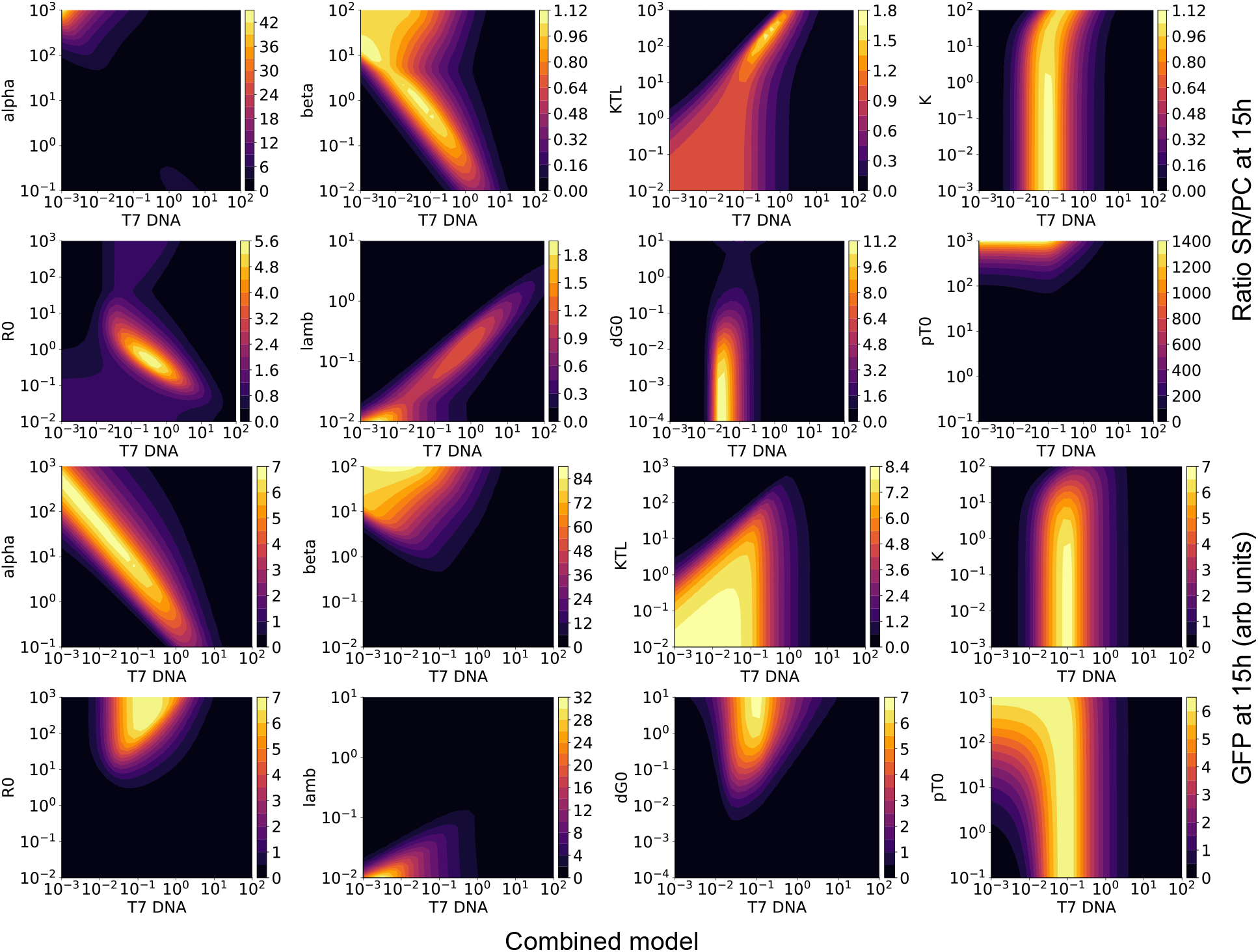
The effects of parameter variations on SR/PC ratio (top) and total eGFP yield (bottom) for the combined model. The more complex model is sensitive to parameter variations, requiring fine-tuning to recapitulate experimental results.

**Figure S19:**
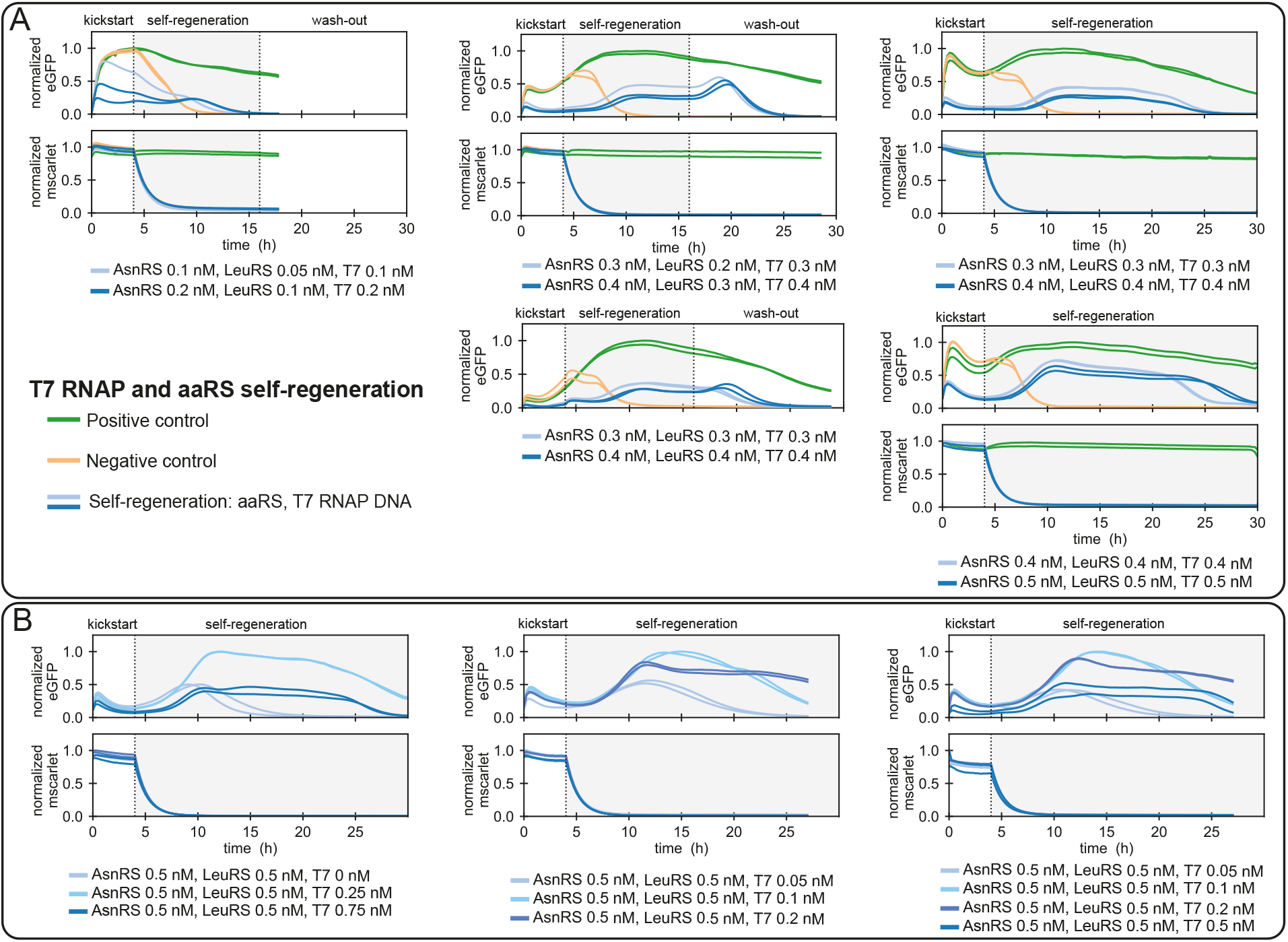
Result summary of regeneration experiments with multiple components being regenerated. Regeneration of AsnRS, LeuRS and T7 RNAP is shown in **(A)**. Titration of T7 RNAP DNA template is depicted in **(B)**. The corresponding mScarlet traces for the given experiments are shown. The level of eGFP intensity is normalised to the maximum intensity obtained in the positive control or to the overall maximum intensity if no positive control was included. PURE composition used for the regeneration experiments are given in Supplementary Table S3. 2 nM of eGFP DNA template was used, and other DNA template concentrations are indicated in the corresponding graphs.

**Figure S20:**
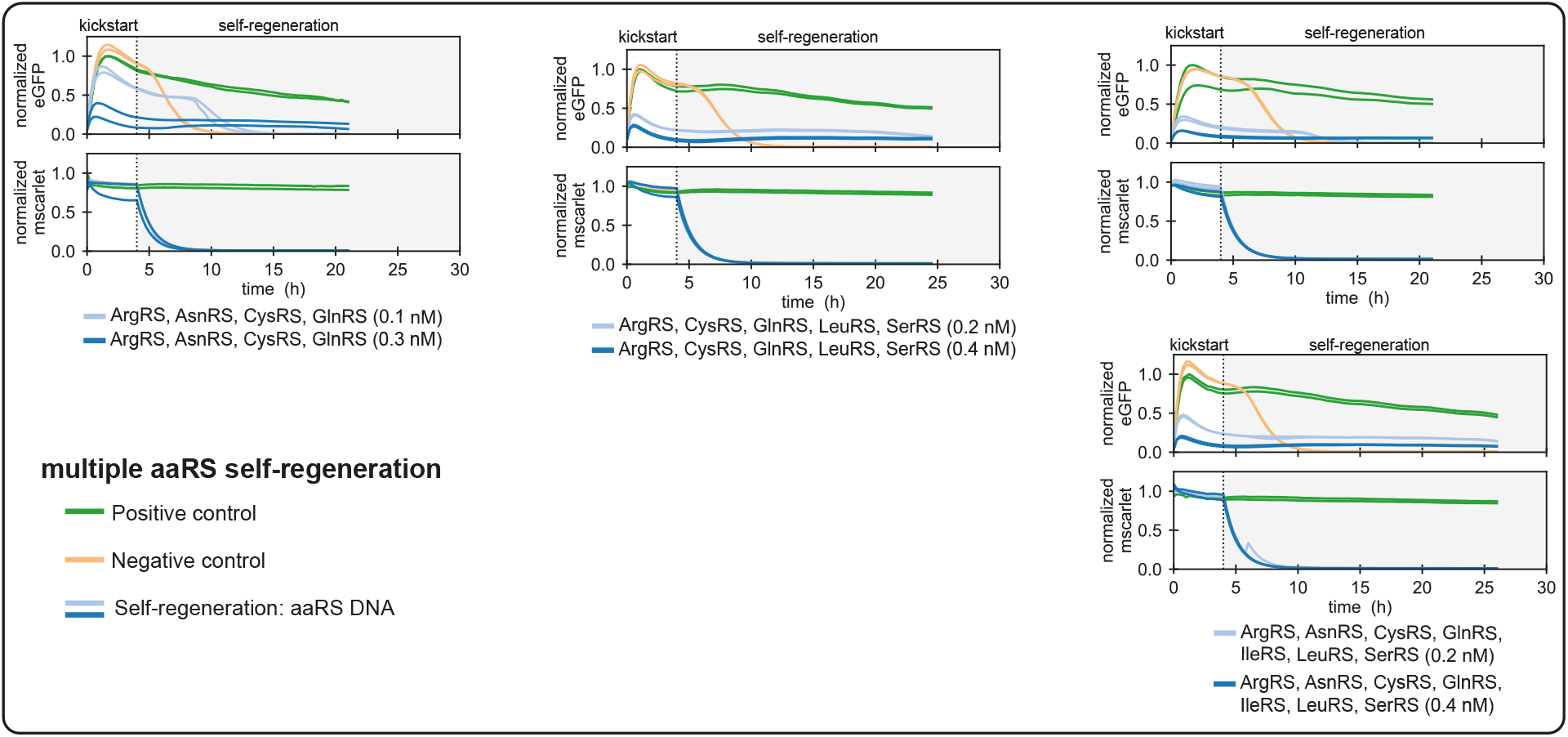
Result summary of multiple aaRSs protein regeneration experiments. Corresponding mScarlet traces for the given experiments are shown. The level of eGFP intensity is normalised to the maximum intensity obtained in the positive control. PURE composition used for the regeneration experiments are given in Supplementary Table S3. 2 nM of eGFP DNA template was used, and other DNA template concentrations are indicated in the corresponding graphs.

**Figure S21:**
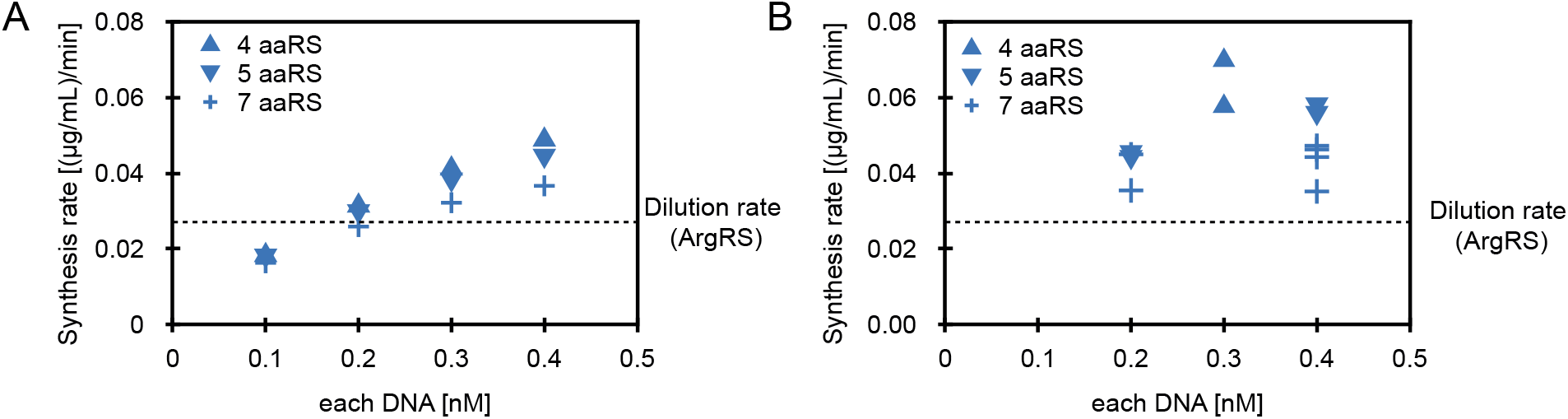
**(A)** Theoretical synthesis rate for single component in multiple components expression, calculated based on eGFP synthesis rate in a microfluidic chemostat (0.44 (*μ*g/mL)/min) and DNA loading in DNA saturated system. **(B)** Estimated synthesis for each component at different DNA concentrations based on the difference in eGFP synthesis rate for positive control and self-regeneration experiment at 15 hours. The eGFP synthesis rate was calculated based on an eGFP calibration curve (Supplementary Fig. S23B) and dilution rate. Dashed line represents the dilution rate of the highest concentrated component (ArgRS), based on the input component concentrations (Supplementary Table S4).

**Figure S22:**
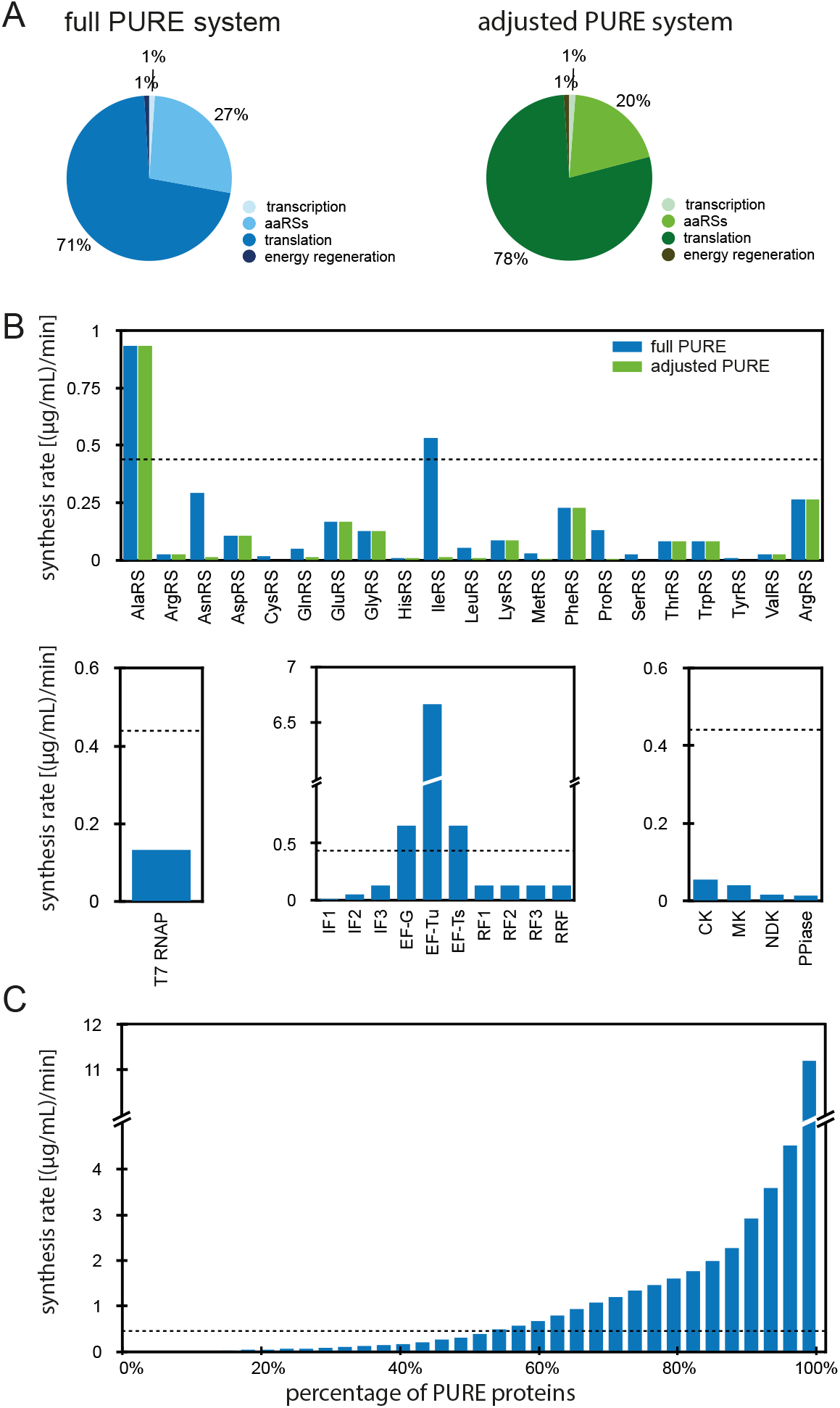
**(A)** Schematic representation of the composition of the full PURE system and adjusted PURE system used for multiple components regeneration. Detailed compositions are given in Supplementary Table S3. **(B)** Estimated minimal required synthesis rate of each PURE component based on dilution rate of each component (Supplementary Table S4) in comparison to the PURE synthesis rate (dashed line). **(C)** Estimated required cumulative synthesis rate for the regeneration of different PURE protein percentage in comparison to the PURE synthesis rate (dashed line). The PURE synthesis rate was calculated based on eGFP expression in a microfluidic chemostat.

**Figure S23:**
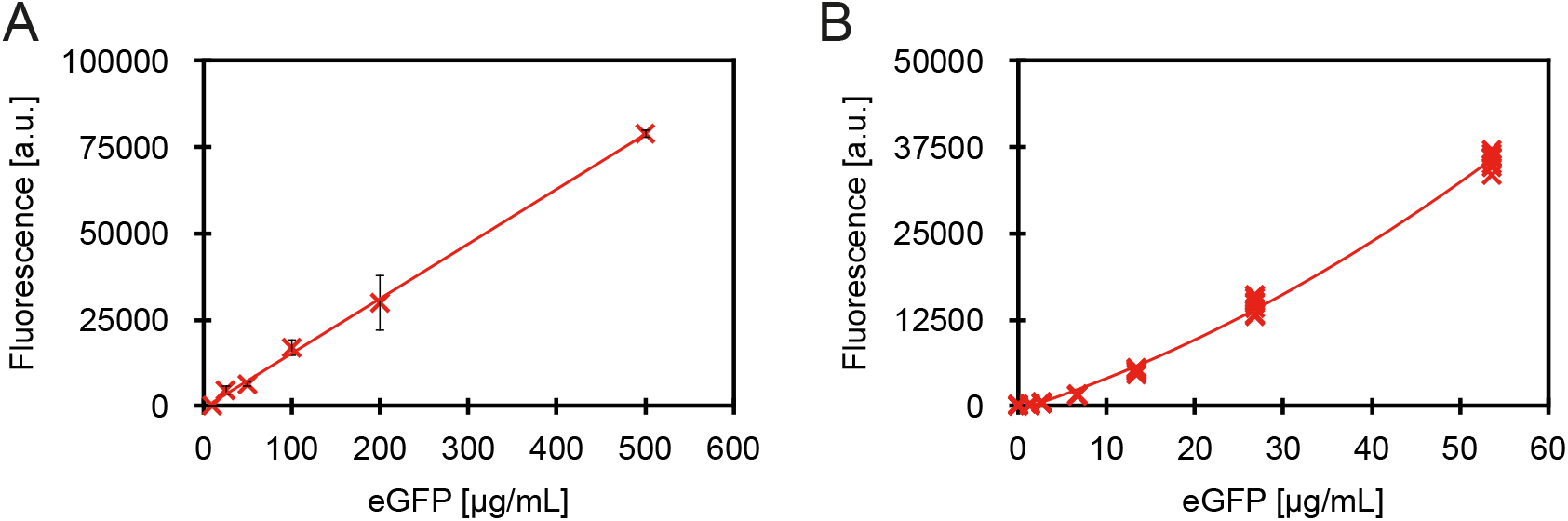
Calibration curve for different eGFP (TP790050, AMS Biotechnology) concentrations in PBS. **(A)** Plate-reader: the standard curve was produced by measuring fluorescence over 60 min with the same settings as for *in vitro* expression. Excitation and emission wavelengths were 488 nm and 507 nm, respectively. Experiments were performed in triplicates. Fluorescence measurements in the first 20 min were not considered. Values are mean ± s.d. (n = 3). **(B)** Microfluidic device: the standard curve was produced by measuring fluorescence over 10 min with the same settings as for *in vitro* expression. Each point represents individual reactor. The fit errors were not propagated as they were negligible compared to experimental errors.

**Table S1:** Microfluidic chip operations for self-regeneration experiments, including positive, and negative controls

| Initial fill |  |  |  |
| --- | --- | --- | --- |
| Step | Operation | Solution | Ring number |
| Repeat the following steps every 15 min |  |  |  |
| **0B Energy solution addition** |  |  |  |
|  | Flush rings | Buffer | 1-8 |
|  | Flush rings | Energy solution | 1-8 |
| **0C PURE solution addition** |  |  |  |
|  | Load 40% (Flush through outlet 60%) | PURE | 1-8 |
| **0D DNA solution addition** |  |  |  |
|  | Load 20% (Flush through outlet 20%) | GFP DNA | 1-4 |
|  | Load 20% (Flush through outlet 20%) | GFP DNA & protein DNA 1 | 5-6 |
|  | Load 20% (Flush through outlet 20%) | GFP DNA & protein DNA 2 | 7-8 |
| **0E Mix** |  |  |  |
| Follow with dilution steps after 15 min |  |  |  |

| kickstart |  |  |  |
| --- | --- | --- | --- |
| Step | Operation | Solution | Ring number |
| Repeat the following steps every 15 min |  |  |  |
| **1A Image each reactor** |  |  |  |
| **Replace 20% of the ring content** |  |  |  |
| **1B Energy solution addition** |  |  |  |
|  | Flush through outlet 20% | Buffer | 1-8 |
|  | Load 8% (Flush through outlet 20%) | Energy solution | 1-8 |
| **1C PURE solution addition** |  |  |  |
|  | Flush through outlet 12% | Buffer | 1-8 |
|  | Load 8% (Flush through outlet 12%) | PURE | 1-8 |
| **1D DNA solution addition** |  |  |  |
|  | Flush through outlet 4% | Buffer | 1-8 |
|  | Load 4% (Flush through outlet 4%) | GFP DNA | 1-4 |
|  | Load 4% (Flush through outlet 4%) | GFP DNA & protein DNA 1 | 5-6 |
|  | Load 4% (Flush through outlet 4%) | GFP DNA & protein DNA 2 | 7-8 |
| **1E Mix** |  |  |  |
| Repeat from the step 1A-E |  |  |  |

| self-regeneration |  |  |  |
| --- | --- | --- | --- |
| Step | Operation | Solution | Ring number |
| Repeat the following steps every 15 min |  |  |  |
| **2A Image each reactor** |  |  |  |
| **Replace 20% of the ring content** |  |  |  |
| **2B Energy solution addition** |  |  |  |
|  | Flush through outlet 20% | Buffer | 1-8 |
|  | Load 8% (Flush through outlet 20%) | Energy solution | 1-8 |
| **2C PURE solution addition** |  |  |  |
|  | Flush through outlet 12% | Buffer | 1-8 |
|  | Load 8% (Flush through outlet 12%) | PURE | 1-2 |
| | Load 8% (Flush through outlet 12%) | ΔPURE | 3-8 |
| **2D DNA solution addition** |  |  |  |
|  | Flush through outlet 4% | Buffer | 1-8 |
|  | Load 4% (Flush through outlet 4%) | GFP DNA | 1-4 |
|  | Load 4% (Flush through outlet 4%) | GFP DNA & protein DNA 1 | 5-6 |
|  | Load 4% (Flush through outlet 4%) | GFP DNA & protein DNA 2 | 7-8 |
| **2E Mix** |  |  |  |
| Repeat from the step 2A-E |  |  |  |

| wash-out |  |  |  |
| --- | --- | --- | --- |
| Step | Operation | Solution | Ring number |
| Repeat the following steps every 15 min |  |  |  |
| **3A Image each reactor** |  |  |  |
| **Replace 20% of the ring content** |  |  |  |
| **3B Energy solution addition** |  |  |  |
|  | Flush through outlet 20% | Buffer | 1-8 |
|  | Load 8% (Flush through outlet 20%) | Energy solution | 1-8 |
| **3C PURE solution addition** |  |  |  |
|  | Flush through outlet 12% | Buffer | 1-8 |
|  | Load 8% (Flush through outlet 12%) | PURE | 1-2 |
| | Load 8% (Flush through outlet 12%) | ΔPURE | 3-8 |
| **3D DNA solution addition** |  |  |  |
|  | Flush through outlet 4% | Buffer | 1-8 |
|  | Load 4% (Flush through outlet 4%) | GFP DNA | 1-8 |
| **3E Mix** |  |  |  |
| Repeat from the step 3A-E |  |  |  |

**Table S2:** Microfluidic chip operations with four self-regeneration experiments

| Initial fill |  |  |  |
| --- | --- | --- | --- |
| Step | Operation | Solution | Ring number |
| Repeat the following steps every 15 min |  |  |  |
| **0B** | **Energy solution addition** |  |  |
|  | Flush rings | Buffer | 1-8 |
|  | Flush rings | Energy solution | 1-8 |
| **0C** | **PURE solution addition** |  |  |
|  | Load 40% (Flush through outlet 60%) | PURE | 1-8 |
| **0D** | **DNA solution addition** |  |  |
|  | Load 4% (Flush through outlet 4%) | GFP DNA & protein DNA 1 | 1-2 |
|  | Load 4% (Flush through outlet 4%) | GFP DNA & protein DNA 2 | 3-4 |
|  | Load 4% (Flush through outlet 4%) | GFP DNA & protein DNA 3 | 5-6 |
|  | Load 4% (Flush through outlet 4%) | GFP DNA & protein DNA 4 | 7-8 |
| **0E** | **Mix** |  |  |
| Follow with dilution steps after 15 min |  |  |  |

| kickstart |  |  |  |
| --- | --- | --- | --- |
| Step | Operation | Solution | Ring number |
| Repeat the following steps every 15 min |  |  |  |
| **1A** | **Image each reactor** |  |  |
| Replace 20% of the ring content |  |  |  |
| **1B** | **Energy solution addition** |  |  |
|  | Flush through outlet 20% | Buffer | 1-8 |
|  | Load 8% (Flush through outlet 20%) | Energy solution | 1-8 |
| **1C** | **PURE solution addition** |  |  |
|  | Flush through outlet 12% | Buffer | 1-8 |
|  | Load 8% (Flush through outlet 12%) | PURE | 1-8 |
| **1D** | **DNA solution addition** |  |  |
|  | Flush through outlet 4% | Buffer | 1-8 |
|  | Load 4% (Flush through outlet 4%) | GFP DNA & protein DNA 1 | 1-2 |
|  | Load 4% (Flush through outlet 4%) | GFP DNA & protein DNA 2 | 3-4 |
|  | Load 4% (Flush through outlet 4%) | GFP DNA & protein DNA 3 | 5-6 |
|  | Load 4% (Flush through outlet 4%) | GFP DNA & protein DNA 4 | 7-8 |
| **1E** | **Mix** |  |  |
| Repeat from the step 1A-E |  |  |  |

| self-regeneration |  |  |  |
| --- | --- | --- | --- |
| Step | Operation | Solution | Ring number |
| Repeat the following steps every 15 min |  |  |  |
| **2A** | **Image each reactor** |  |  |
| Replace 20% of the ring content |  |  |  |
| **2B** | **Energy solution addition** |  |  |
|  | Flush through outlet 20% | Buffer | 1-8 |
|  | Load 8% (Flush through outlet 20%) | Energy solution | 1-8 |
| **2C** | **PURE solution addition** |  |  |
|  | Flush through outlet 12% | Buffer | 1-8 |
| | Load 8% (Flush through outlet 12%) | ΔPURE | 1-2 |
| **2D** | **DNA solution addition** |  |  |
|  | Flush through outlet 4% | Buffer | 1-8 |
|  | Load 4% (Flush through outlet 4%) | GFP DNA & protein DNA 1 | 1-2 |
|  | Load 4% (Flush through outlet 4%) | GFP DNA & protein DNA 2 | 3-4 |
|  | Load 4% (Flush through outlet 4%) | GFP DNA & protein DNA 3 | 5-6 |
|  | Load 4% (Flush through outlet 4%) | GFP DNA & protein DNA 4 | 7-8 |
| **2E** | **Mix** |  |  |
| Repeat from the step 2A-E |  |  |  |

**Table S3:**
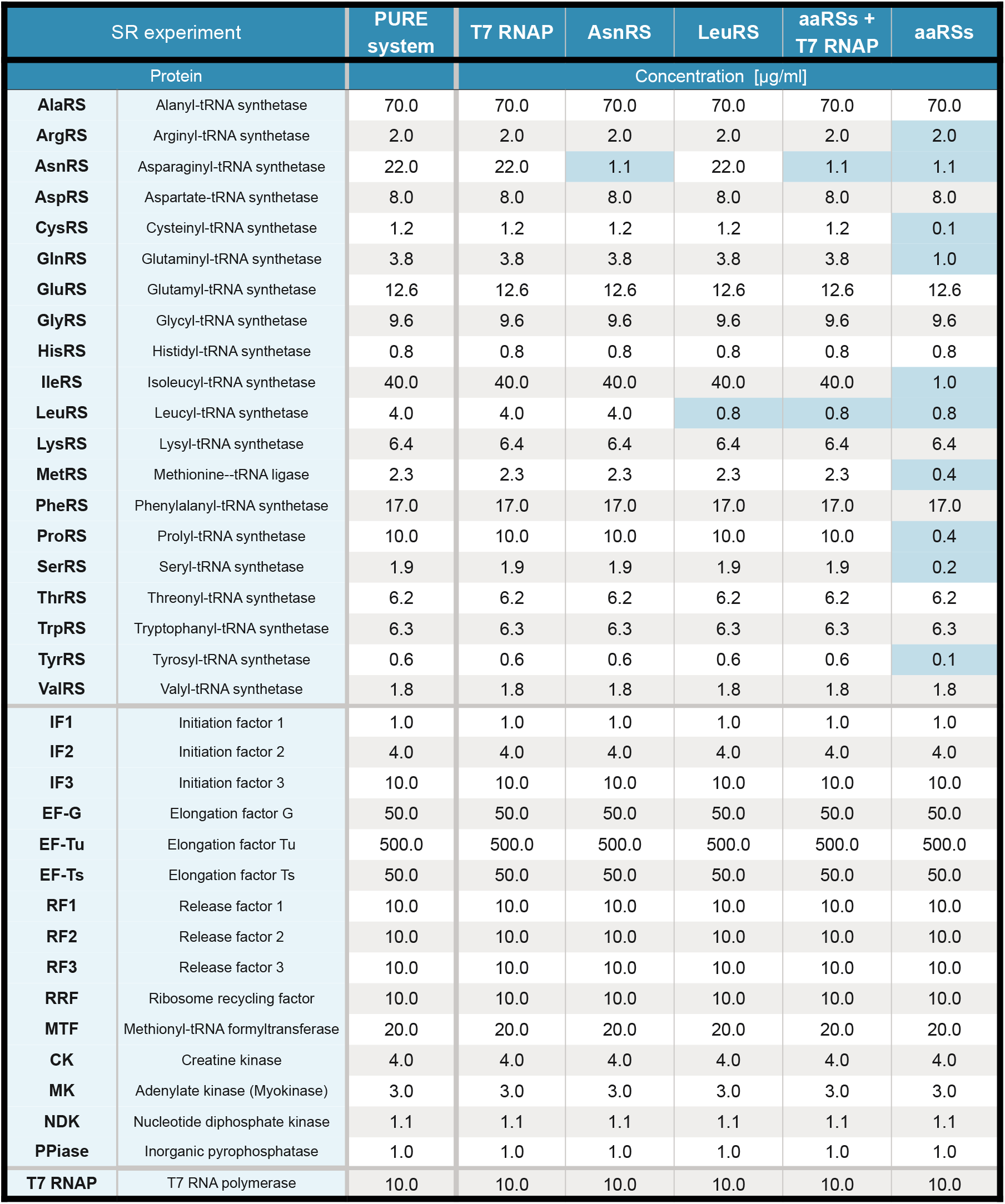
PURE system formulations used

**Table S4:**
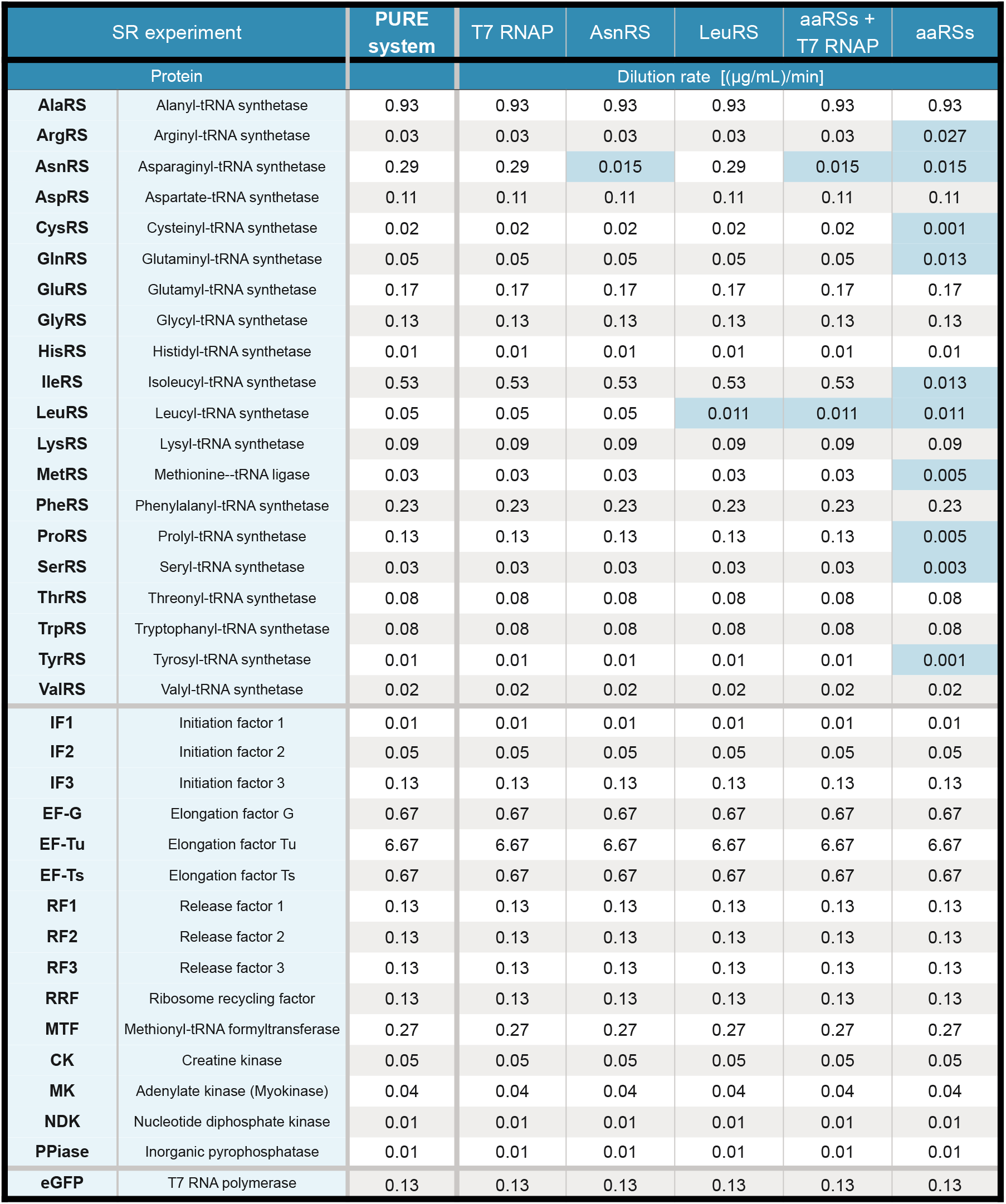
Calculated dilution rates based on concentrations in Table S3

**Table S5:** Replenishing schedule for modeling the three-stage experiment

| Self-regeneration | Species replenished |
| --- | --- |
| Stage 1 | $R, d_T, d_G, p_T$ |
| Stage 2 | $R, d_T, d_G$ |
| Stage 3 | $R, d_G$ |
| Positive control | Species replenished |
| Stage 1 | $R, d_G, p_T$ |
| Stage 2 | $R, d_G, p_T$ |
| Stage 3 | $R, d_G, p_T$ |

**Table S6:**
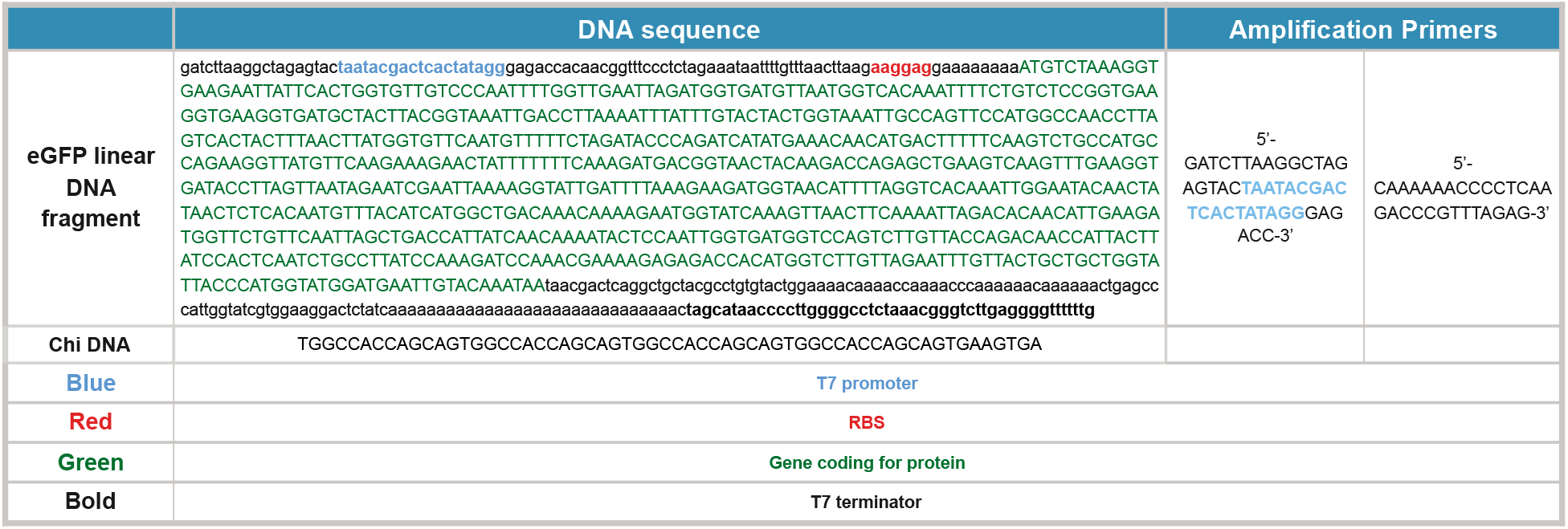
DNA sequences

**Table S7:**
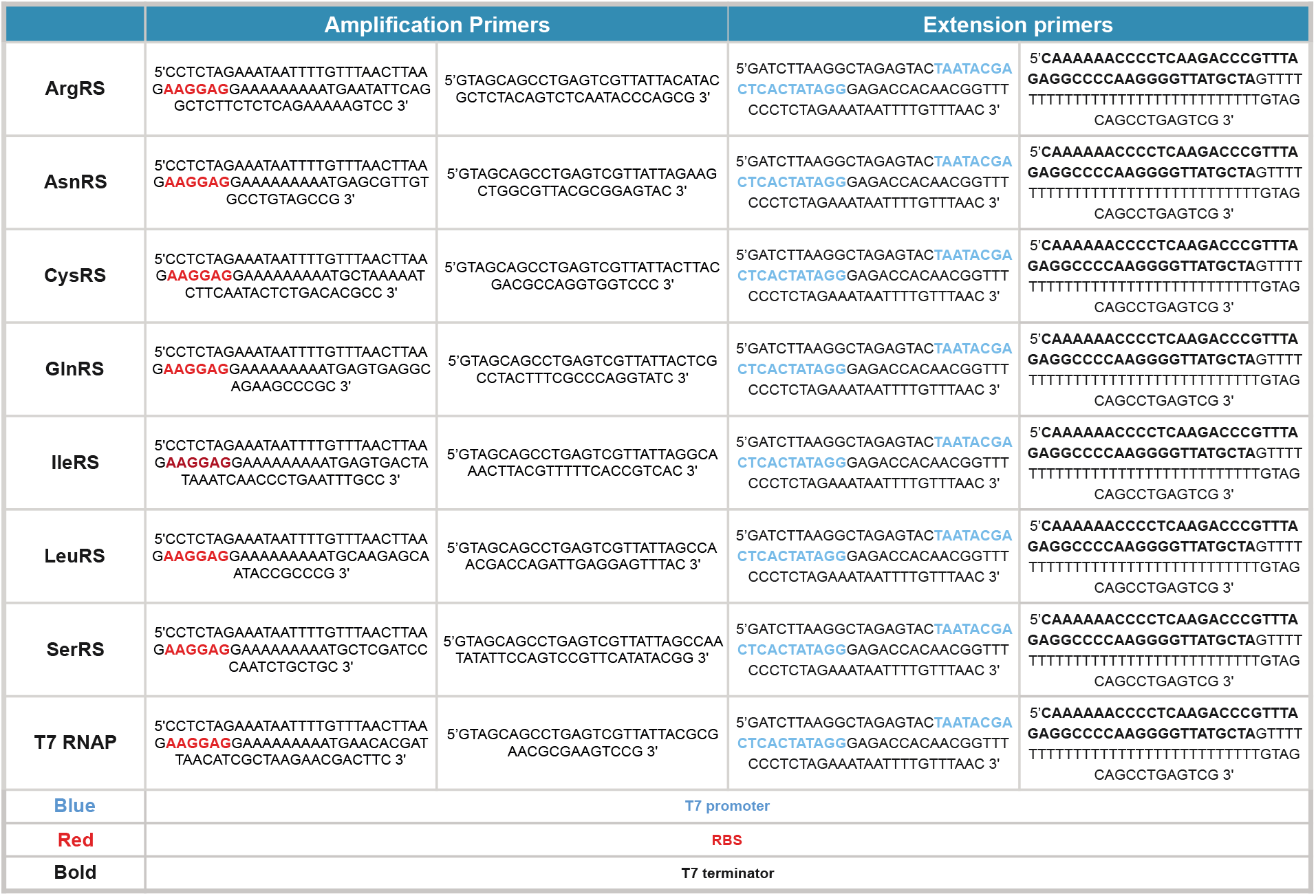
Primer sequences

**Table S8:**
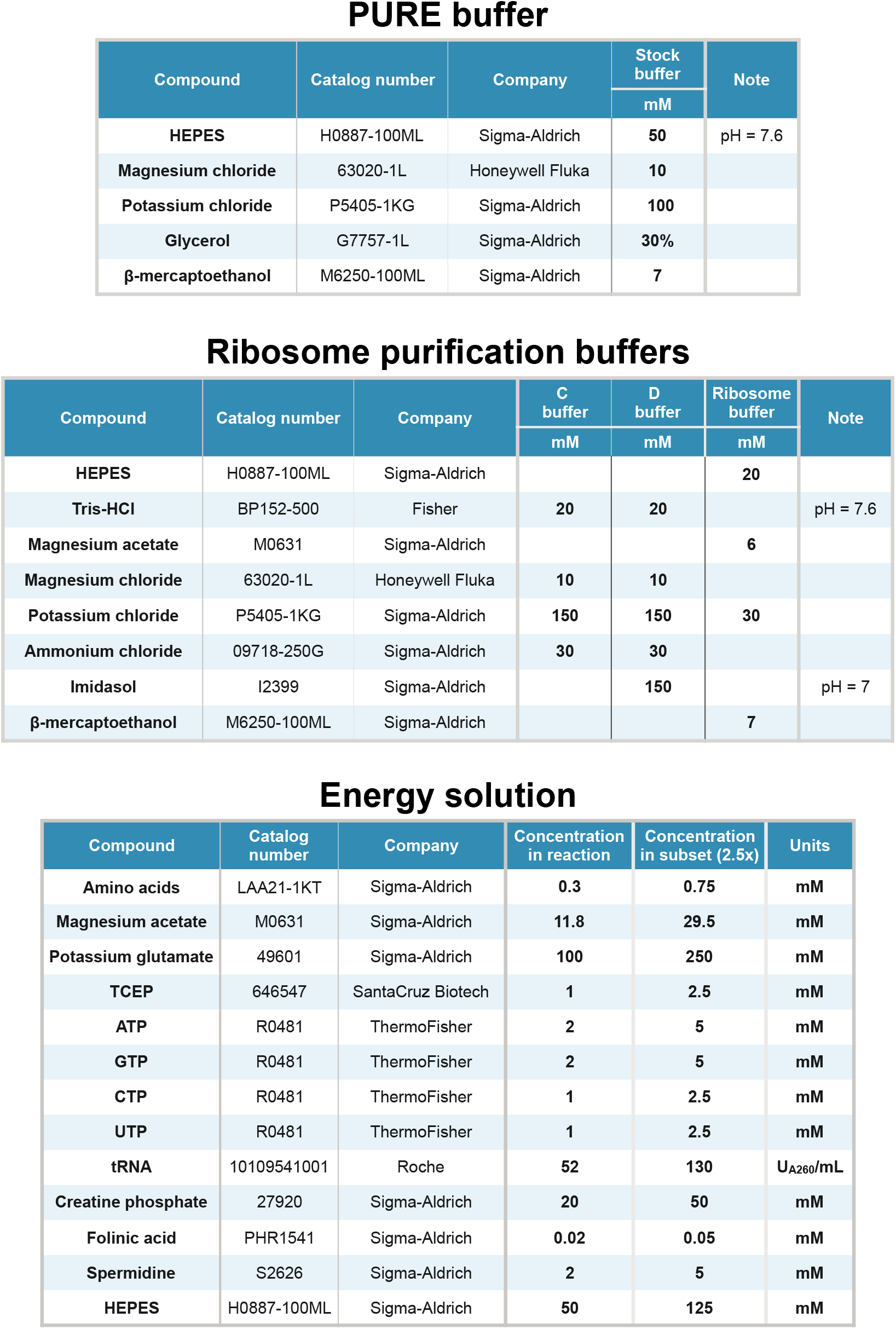
Buffers and energy solution

